# RNA Folding Nearest Neighbor Parameters Including the Modification 1-Methyl-Pseudouridine

**DOI:** 10.64898/2026.04.09.717343

**Authors:** Elzbieta Kierzek, Thandolwethu S. Shabangu, Olivia M. Hiltke, Megan Miaro, Sebastian Arteaga, Brent M. Znosko, Elizabeth A. Jolley, Philip C. Bevilacqua, John SantaLucia, Holly A. SantaLucia, Haining Lin, Mihir Metkar, Sharon Aviran, Marta Soszyńska-Jóźwiak, Ryszard Kierzek, David H. Mathews

## Abstract

Nearest neighbor analysis is commonly used to estimate RNA folding stabilities. In this contribution, we report a set of RNA folding nearest neighbor parameters for estimating free energy change for RNA sequences including 1-methyl-pseudouridine. Development of mRNA vaccines has identified 1-methyl-pseudouridine as a key nucleobase modification for suppressing innate immune responses. However, the contributions of these modifications to RNA folding stability were unclear. Our new parameters provide helical terms for 1-methyl-pseudouridine-adenine and 1-methyl-pseudouridine-guanine base pairs. The parameters also estimate loop stabilities for loops with 1-methyl-pseudouridine or a combination of 1-methyl-pseudouridine and uridine. These parameters are derived using 208 optical melting experiments and tested against an additional 16 optical melting experiments. On average, we find that substitution of uridine with 1-methyl-pseudouridine stabilizes RNA folding, with the extent of stabilization depending on adjacent sequence. The estimation of tRNA folding ensembles for tRNA sequences with 1-methyl-pseudouridine was significantly improved using the new nearest neighbor parameters. The new nearest neighbor parameters are provided as part of the RNAstructure software package. With these parameters, the secondary structures of natural sequences with 1-methyl-pseudouridine and mRNA therapeutics fully substituted with 1-methyl-pseudouridine can be modeled.

## Introduction

RNA is a biological macromolecule that is appreciated for its roles in protein expression (mRNA, rRNA, and tRNA), as a catalyst, as a regulator of gene expression, and as an adapter for molecular recognition^1, 2^. It is also an important drug target and therapeutic.

It is increasingly appreciated that RNA sequences use an alphabet of nucleotides beyond the canonical adenosine (A), cytidine (C), guanosine (G), and uridine (U). Covalent modifications of these nucleotides occur on both the base and sugar-phosphate backbone and are found naturally or produced synthetically. It has long been known that tRNA nucleotide modifications are needed for tRNA function, and that modifications are conserved in snRNA and rRNA^3^. With the development of next generation sequencing methods that can identify modified nucleotides, we now appreciate the wide usage of modified nucleotides in coding and non-coding mRNA. A database catalogs the known modifications^4^.

One important RNA base modification is 1-methyl-pseudouridine (1mΨ; Figure 1A). Pseudouridine (Ψ) is an isomer of uridine in which the glycosidic bond to the N1 base position is replaced with a glycosidic bond to the C5 position^5^. In 1mΨ, the N1 is methylated. This modification occurs naturally in 18S rRNA and archaeal tRNA^6^. 1mΨ is also widely substituted for U in vaccine and therapeutic mRNAs to evade the innate immune response^7^.

**Figure 1.**
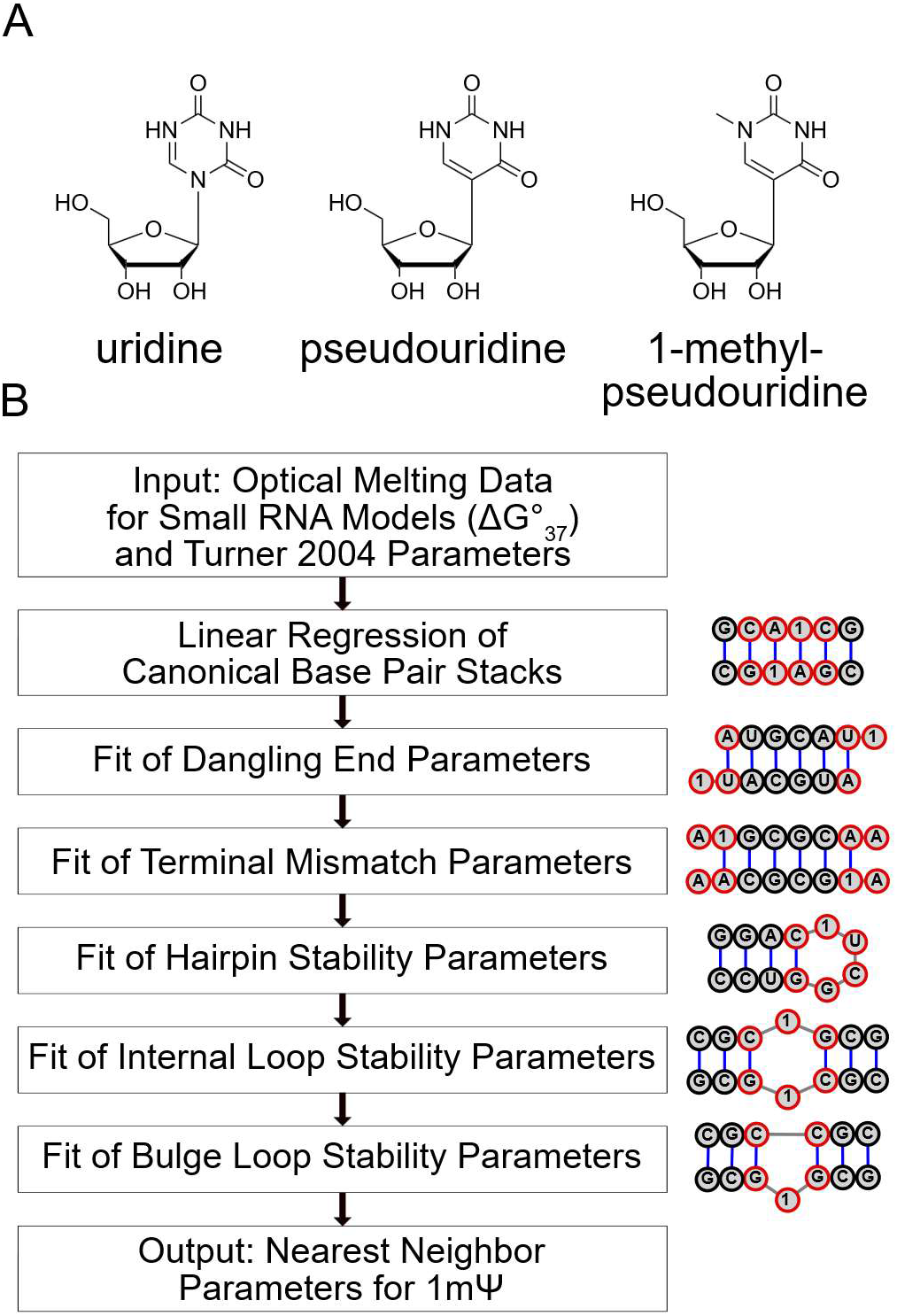
Overview and parameter fitting procedure. Panel A shows uridine, pseudouridine (Ψ), and 1-methyl-pseudouridine (1mΨ). Panel B shows the fitting procedure used to determine folding free energy parameters for sequences that include 1mΨ. The input is a set of optical melting data that provide folding free energy changes for small model systems and the Turner 2004 nearest neighbor parameters. In turn, the base pair stack, dangling end, terminal mismatch, hairpin loop, internal loop, and bulge loop terms are fit for 1mΨ terms. The output is the set of nearest neighbor parameters that uses the Turner 2004 functional form and has terms that include 1mΨ. The right column shows a representative model system for each of the motifs being fit, with the motif highlighted in red, where 1 is used to represent 1mΨ.

Often the structure of RNA is important for function. This can be modeled as secondary structure, i.e. the set of the canonical base pairs and the unpaired regions, called loops, that are defined by the pairs^8^. The secondary structure can be predicted from sequence, and often this is done by predicting the lowest free energy structure, which is the most populated structure at equilibrium, or by predicting the base pairing probabilities in the Boltzmann ensemble of structures^9, 10^. To quantify the stability of folding from the random coil to a specific structure, a set of thermodynamic nearest neighbor parameters (a.k.a. Turner rules) can be used to estimate the Gibbs free energy change (ΔG°_37_)^11^. A structure with lower folding free energy change is more stable and thus that structure is more likely to be found at equilibrium than a structure with higher folding free energy change.

The Turner rules were developed to estimate experimentally measured stabilities from optical melting experiments for RNA sequences^12^. The most recent set of parameters, called “Turner 2004,” has terms for helices^13, 14^ and loops^15^. This functional form has been used as the basis for parameter sets that were optimized for RNA structure prediction^16, 17, 18^. Additional features have also been explored by expanding the functional form^19^.

We previously developed a set of nearest neighbor parameters for predicting structures with the bases 6-methyl-adenosine (m^6^A), A, C, G, and U^20^. That work used a set of 45 optical melting experiments, chosen to study the most important sequence features as determined by sensitivity analyses performed by Zuber et al^21, 22^. In a subsequent test of the m^6^A parameters, an additional set of 98 duplexes were studied with either A or m^6^A in the context of the consensus site for methylation in mammals, RRACH (where R is G or A and H is A, C, or G.), ^23^. That work demonstrated that the m^6^A parameters were approximately as accurate as parameters for A in predicting the folding stability of oligonucleotides.

In another study, nearest neighbor parameters were fit to include synthetic DNA base pairs, P-Z, for an alphabet of bases of P (2-amino-8-(1′-b-D-2′-deoxyribofuranosyl)-imidazo-[1,2a]-1,3,5-triazin-[8H]-4-one), Z (6-amino-3-(1′-b-D-2′-deoxyribofuranosyl)-5-nitro-1H-pyridin-2-one), A, C, G, and T^24^. That work also considered G-Z pairs, which were roughly as stable as A-T base pairs. Using these parameters, sequences were designed to fold into challenging secondary structures^25^. By including the synthetic base pairs, the designs were improved by *in silico* evaluation as compared to using the canonical DNA nucleotides alone.

Here, we report nearest neighbor parameters for 1mΨ, A, C, G, and U. Prior work demonstrated that 1mΨ substitution for U can enhance folding stability and our new parameters quantify this^26, 27^. It was challenging to develop these parameters because 1mΨ can form canonical base pairs with A or with G, analogous to A-U and G-U base pairs. This means there are 34 helical parameters to fit, which requires an extensive set of experimental data. It was also important to develop parameters for unpaired nucleotides, as 1mΨ substitution for U can have a large effect on loop stability. A set of 224 optical melting experiments were performed, of which 208 were used for fitting parameters and 16 were used to evaluate the parameters. Figure 1B shows the overall fitting procedure. The Turner 2004 nearest neighbor parameters were used as the basis for the new parameterization because of their widespread use. We demonstrate that the parameters improve the modeling of secondary structures of tRNA sequences that contain 1mΨ. The new parameters were incorporated into the RNAstructure software package and are freely available^28^.

## Results

### Integration of Data from Two Sources and Reproducibility of Optical Melting

To perform the experiments required to fit the nearest neighbor parameters, we integrated optical melting data for oligonucleotides with 1mΨ from two sites, DNA Software, Inc., and the Institute of Bioorganic Chemistry of the Polish Academy of Sciences. Additionally, we relied on comparisons to optical melting studies performed previously with canonical RNA oligonucleotides that established the Turner 2004 nearest neighbor parameters.

To test the reproducibility of optical melting experiments, we chose a set of three calibration duplexes that had previously been studied and that we repeated at DNA Software, Inc., the Institute of Bioorganic Chemistry, and Saint Louis University. Figure 2 provides the sequences and measured ΔG°_37_s and Table S1 provides the full thermodynamic parameters. These sequences were chosen to cover melting temperatures (for 0.1 mM strand concentrations) from 40 to 60 °C. They include self-complementary and non-self-complementary sequences and one sequence with a 2×2 internal loop.

**Figure 2.**
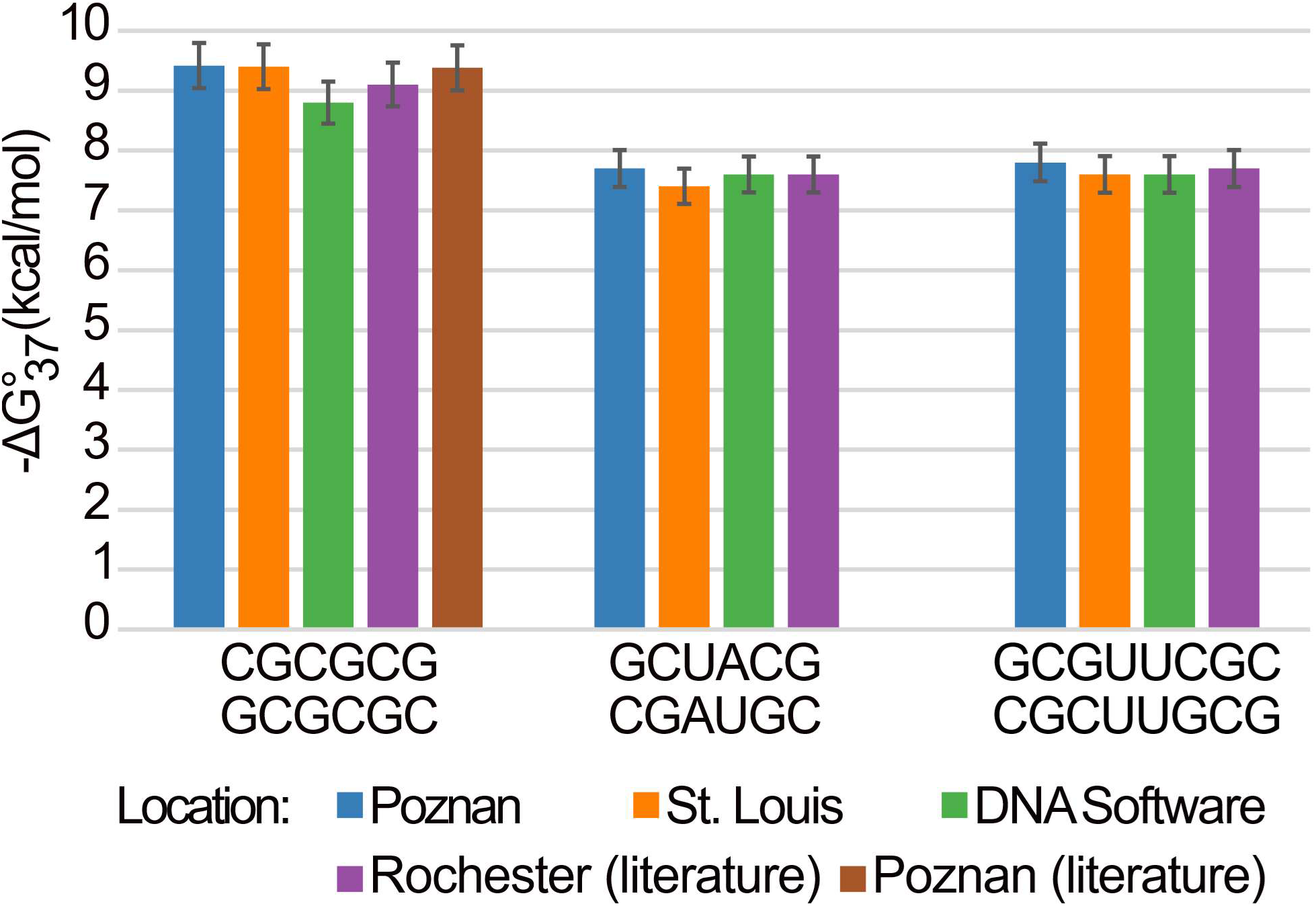
The comparison of optical melting results across sites. Three duplexes were studied by optical melting across three sites (Institute of Bioorganic Chemistry, Poznan; Saint Louis University, St. Louis; and DNA Software). The results were also compared to previous literature results (University of Rochester, Rochester, and Poznan)^13, 20, 64, 65^. Uncertainties are estimated as 4% of the value^13^.

It was previously found that ΔG°_37_ for folding determined by optical melting studies for three DNA sequences studied each at two sites were reproducible within 3% ^29^. On average, for these three RNA duplexes, the uncertainty in ΔG°_37_ was 2.1% (Supplementary Table S1), where the uncertainty is the ratio of the standard deviation of the measured ΔG°_37_ and the absolute value of the mean of the measured ΔG°_37_. This close agreement between sites required temperature calibration for the instruments (Methods). We concluded that, given this agreement, we can integrate folding free energies from multiple sites to fit nearest neighbor parameters.

### Nearest Neighbor Parameter Fit Procedure

Nearest neighbor parameters were fit to the Turner 2004 functional form, including an update for G-U pair helical stacks by Chen et al.^30^. Figure 1 shows the procedure, which follows a specific path so that the required dependencies for analyzing the optical melting data are available. For example, dangling end parameter fits rely on estimates for helical stack parameters, therefore the helical stacks are fit before dangling ends. In turn, hairpin loop fits require estimates for helical stacks and terminal mismatches. The Methods detail each of the fits and parameters that were used.

### Helical parameters

There are 34 nearest neighbor parameters for helical stacks that involve 1mΨ in base pairs with A or G. These were fit using a set of 81 model duplexes composed of fully base paired helices (Table S2). In the fit, the helical stacks with two canonical base pairs were fixed with their previously determined values^13, 30^. The Watson-Crick-Franklin stacks are those used in the Turner 2004 parameters^11^, but the GU stacks are from a more recent compilation^30^.

Figure 3 and Table S3 compare the stacks involving 1mΨ-A pairs to the analogous stacks including Watson-Crick-Franklin pairs. For these pairs, each substitution of 1mΨ for U stabilizes the ΔG°_37_ by an average of −0.32 ± 0.17 kcal/mol, although this varies from −0.56 to −0.14 kcal/mol depending on the sequence context. Figure 4 and Table S4 compare the stacks involving 1mΨ-G pairs or 1mΨ-A adjacent to U-G pairs to the analogous canonical base pairs. For these base pair stacks, each substitution of 1mΨ for U stabilizes by a similar average of −0.27 ± 0.48 kcal/mol, with an even wider range from −1.43 to 0.73 kcal/mol.

**Figure 3.**
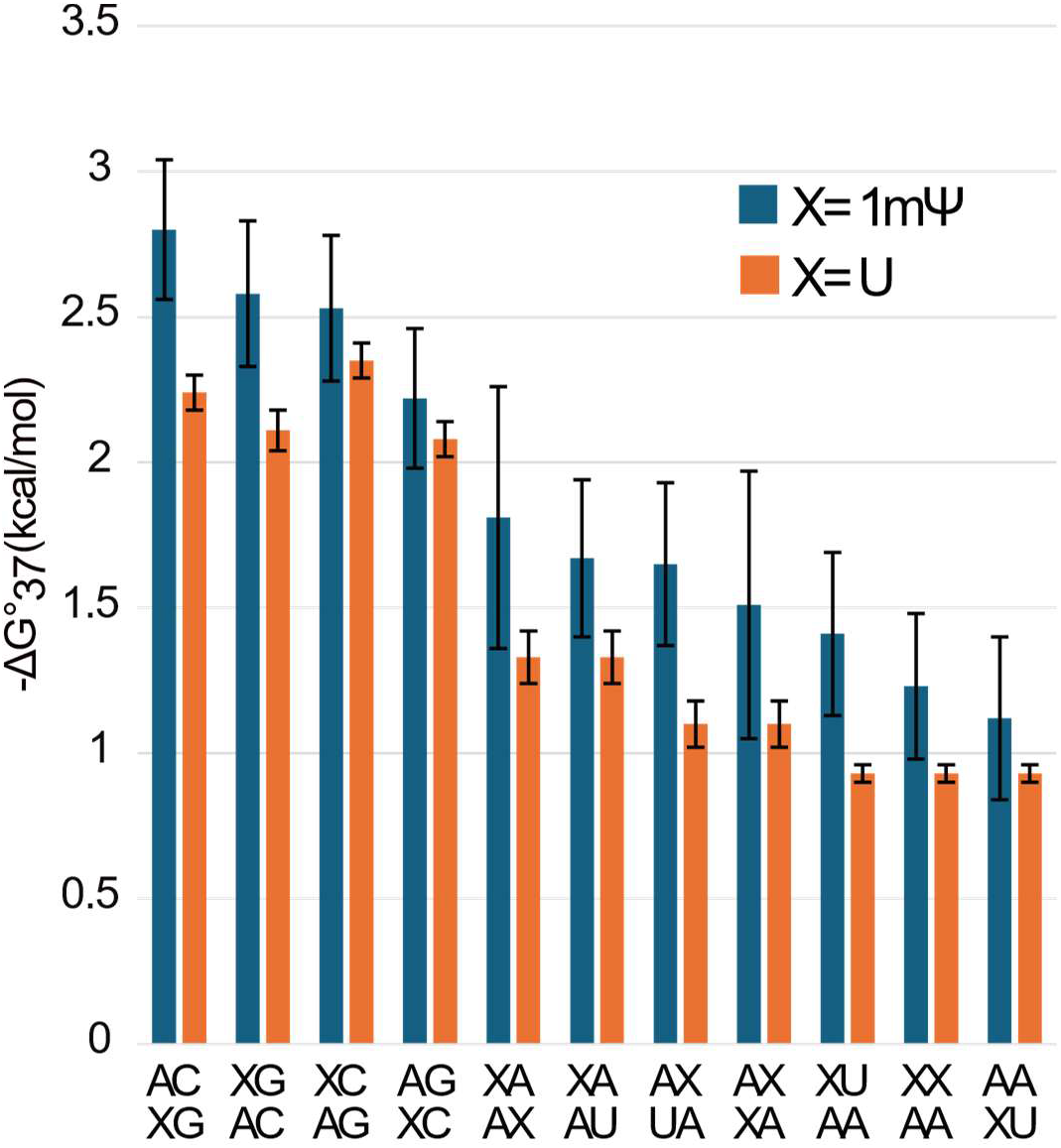
Comparison of 1mΨ-A base pair stacking parameters to analogous U-A base pair parameters. On average, each 1mΨ-A substitution for a U-A is stabilizing by −0.32 ± 0.17 kcal/mol.

**Figure 4.**
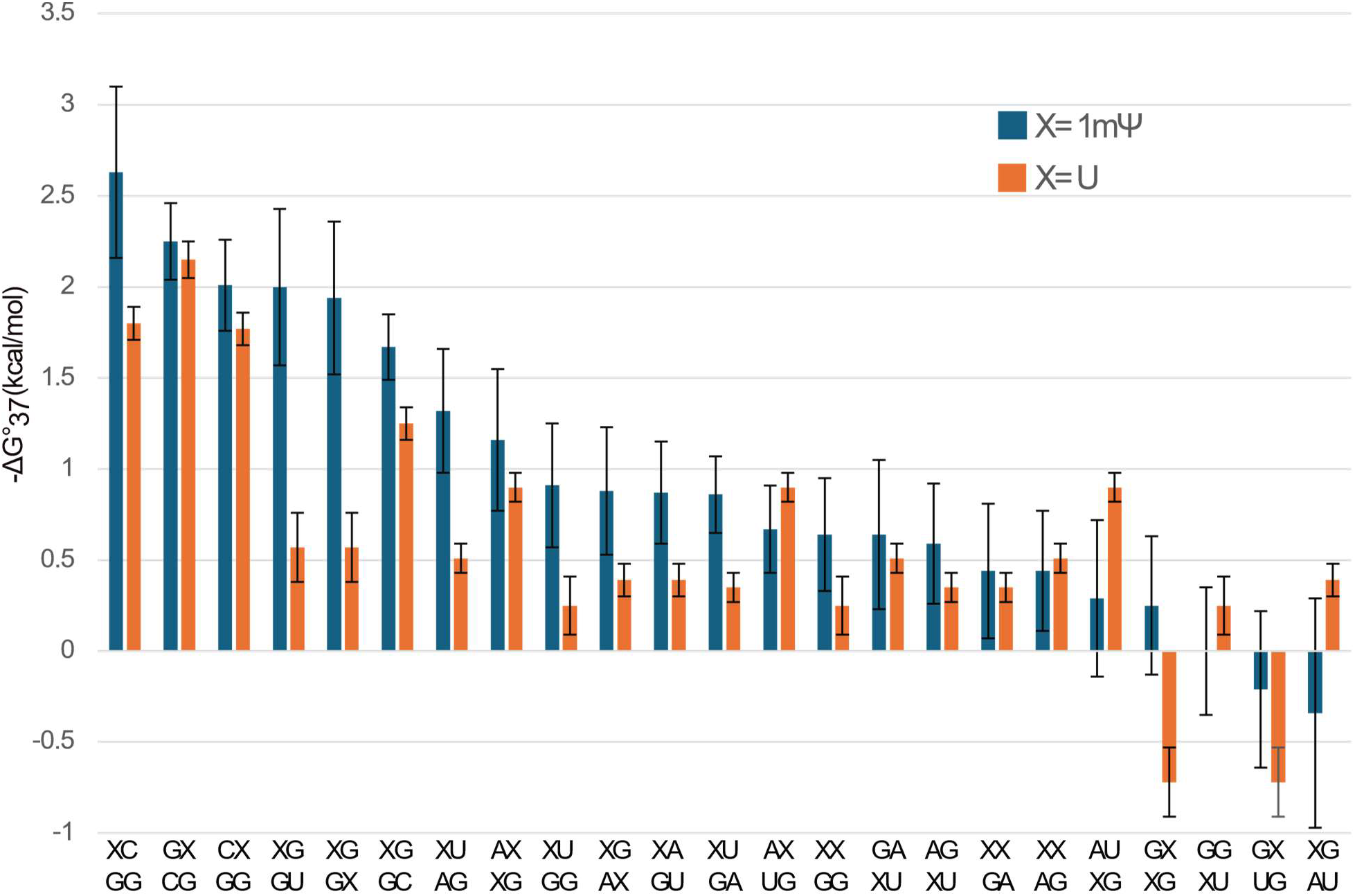
Comparison of 1mΨ base pair stacking parameters to analogous U base pair parameters for stacks with 1mΨ-G or U-G pairs. On average, each 1mΨ-G substitution for a U-G is stabilizing by −0.27 ± 0.48 kcal/mol.

Interestingly, we found that terminal 1mΨ-A and 1mΨ-G pairs require no stability penalty. In the Turner 2004 rules, terminal U-A pairs destabilize a helix by 0.45 kcal/mol^13^. Terminal U-G pairs, however, also do not destabilize a helix^30^.

The nearest neighbor parameters agree well with the experiments. Figure 5 and Table S5 show the estimated helical stability as a function of measured helical stability for all duplexes. The 81 duplexes used to fit the nearest neighbor parameters were measured at the Institute of Bioorganic Chemistry. These have a root mean squared deviation (RMSD) of 0.40 kcal/mol between the measured stability and the estimated stability. An additional twelve 1mΨ-containing duplexes were measured at DNA Software (Table S6). These have stabilities also well estimated by the nearest neighbor parameters, with the exception of two outliers that differ from experimental values by over three standard errors. The overall RMSD for the DNA Software duplexes is 1.23 kcal/mol, but this is reduced to 0.68 kcal/mol when the two outliers are removed.

**Figure 5.**
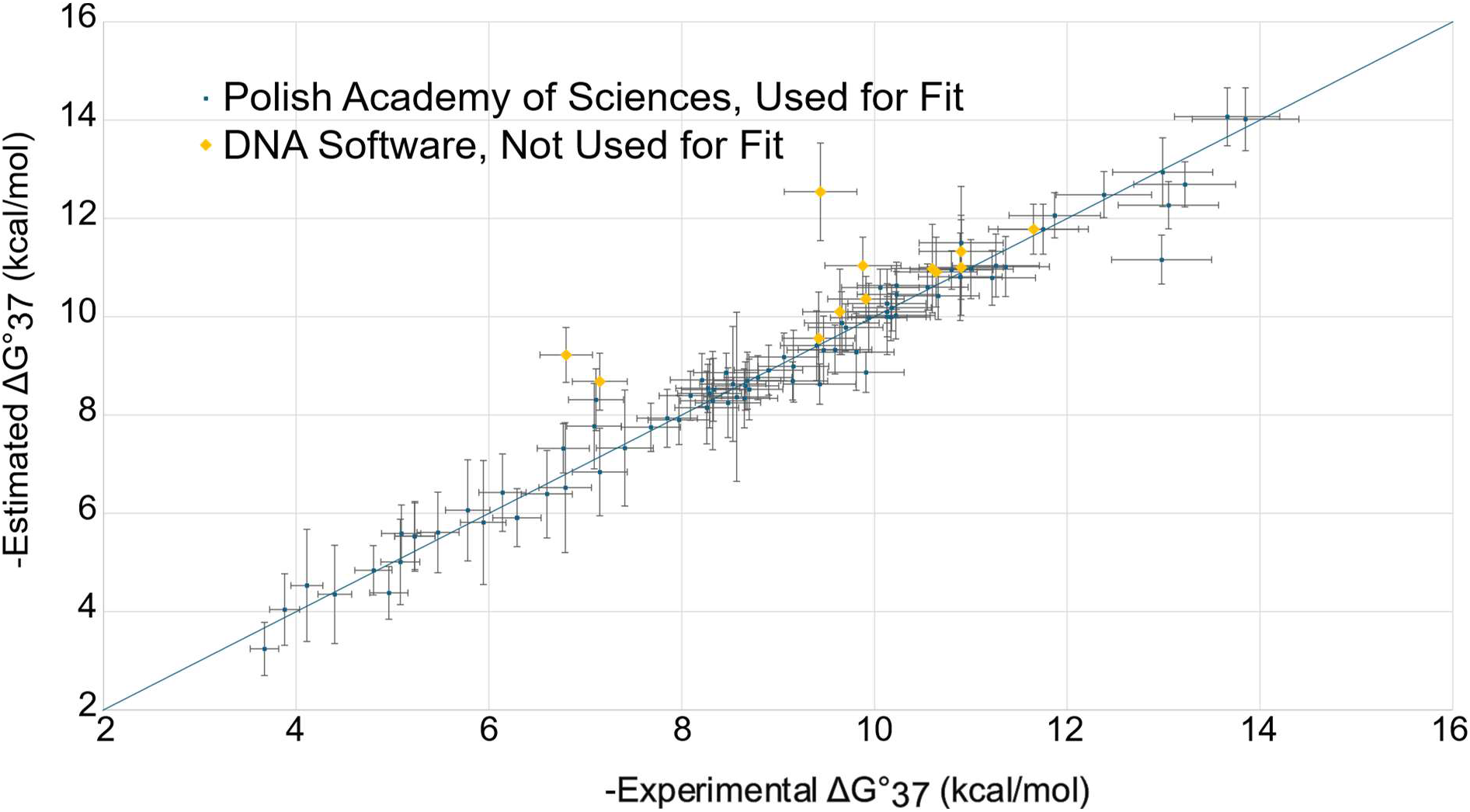
Agreement between experimentally measured helical free energy changes and nearest-neighbor estimated free energy changes. The dark squares represent helices measured at the Institute of Bioorganic Chemistry and used to fit the parameters. The light diamonds are helices measured by DNA Software and not used for the fits. The line is the diagonal, which represents perfect agreement between estimations and experiments. The experimental uncertainties are 4% of the values, estimating reproducibility across sites^13^. The uncertainties in the estimates are from propagation of standard errors of the regression from individual terms.

We find the uncertainties for the helical parameters with 1mΨ are larger in magnitude than those for analogous stacks with U (Figures 3 and 4 and Tables S4 and S5). In general for linear regression when the data fit the model, the uncertainty for parameters tends to decrease with increasing data. For the 1mΨ parameters, we used 81 experiments for the fit; for U there were 90 helices used to fit the Watson-Crick-Franklin stacks and an additional 70 helices with G-U pairs. Those experiments with U were collected over the time period from 1981 to 2012^30, 31^. If more data became available for 1mΨ, this model could be refined.

### Terminal Mismatches and Dangling Ends

Dangling ends, which are a single unpaired nucleotide stack on a helix end, and terminal mismatches, which are non-canonical pairs at a helix end, both stabilize helix formation (Fig. 1). Terms that represent these effects in the nearest neighbor model are sequence dependent. We studied 20 dangling end-models (Table S7) and 40 terminal mismatch models (Table S8) by optical melting. The motif stabilities for dangling ends and terminal mismatches are in Table S9 and S10, respectively.

For dangling ends, the most striking effect is that 3’ dangling ends are more stabilizing when adjacent to a 1mΨ-A pair than when adjacent to an analogous U-A pair. For example, the A, C, and G 3’ dangling ends on average are more stable by −0.43 ± 0.02 kcal/mol. A 5’ dangling 1mΨ is slightly more stabilizing than an analogous U on AU, CG, and GC helix ends by −0.22 ± 0.07 kcal/mol. A 3’ dangling 1mΨ, however, is apparently slightly less stabilizing on average than an analogous U on these helix ends by 0.10 ± 0.07 kcal/mol.

Terminal mismatches are stabilized when they contain 1mΨ as compared to an analogous U. For example, 1mΨ-1mΨ mismatches adjacent to A-1mΨ and 1mΨ-A terminal pairs are more stabilizing than analogous motifs by −0.39 ± 0.41 and −0.88 ± 0.58 kcal/mol, respectively. 1mΨ-1mΨ and 1mΨ-C mismatches adjacent to C-G and G-C terminal pairs are stabilizing on average by −0.46 ± 0.14 kcal/mol.

### Hairpin Loops

Eight hairpin stem-loops containing 1mΨ were studied by optical melting, and an additional four were studied with an analogous U to quantify the stability changes from 1mΨ substitution (Table S11). In the Turner 2004 model^15^, hairpin loop stabilities are estimated with a model that considers the number of unpaired nucleotides and the sequence of the closing base pair and first mismatch, i.e. the nucleotides adjacent to the closing base pair. The sequence-dependent term is the terminal mismatch term with additional stability added for UU, GA, or GG first mismatches. Additionally, a subset of triloops (hairpins with three unpaired nucleotides), tetraloops (four unpaired nucleotides), and hexaloops (six unpaired nucleotides) have stabilities that are poorly modeled by the generic model^15^. These sequences have folding free energy increments that are in lookup tables. The eight hairpins with 1mΨ were designed to test whether the Turner 2004 model can be used by placing the 1mΨ in the closing pair, in the first mismatch, in an unpaired position or helical position away from the first mismatch, and in a special tetraloop hairpin.

The Turner 2004 model for hairpin loops works well for modeling the stability of hairpin loops containing 1mΨ (Figure 6 and Table S12)^15^. The term in the model that additionally stabilizes U-U first mismatches provides an additional −0.95 ± 0.81 kcal/mol stability beyond the U-U terminal mismatch value^15, 22^. We found that 1mΨ-U first mismatches are also more stable than the 1mΨ-U terminal mismatches alone would suggest (average of −0.78 ± 1.18 kcal/mol for two sequences), and therefore we apply the same −0.95 kcal/mol stabilization to U-U, U-1mΨ, 1mΨ-U, and 1mΨ −1mΨ first mismatches.

**Figure 6.**
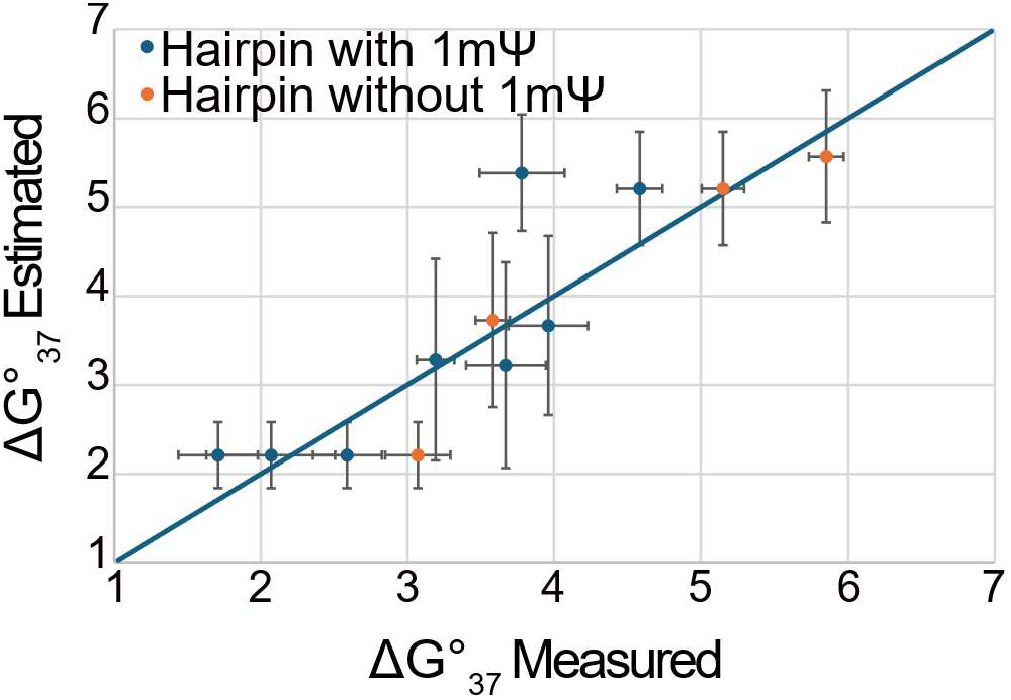
Hairpin loop stabilities are well modeled using the Turner 2004 functional form. This shows the nearest-neighbor-estimated hairpin loop stabilities as a function of measured hairpin loop stabilities (Table S12). The diagonal line is shown. The blue points are for hairpin loops that included 1mΨ. The control experiments, with U instead of 1mΨ are shown for reference in blue.

We found that the stabilities of hairpins C(1mΨ)UCGG and CU(1mΨ)CGG are well described by the lookup value of the tetraloop sequence CUUCGG^15, 32, 33^, with differences of −0.5 ± 0.46 and 0.38 ± 0.44 kcal/mol, respectively. Based on these results, where the differences in stability between tetraloops with 1mΨ or U in analogous positions are within the experimental uncertainty, we apply the lookup values for triloops, tetraloops, and hexaloops for all variations of the sequences where U is replaced with 1mΨ.

### Internal Loops

We studied 47 model systems with internal loops by optical melting (Table S13). These are interruptions in helices with unpaired nucleotides on each side of the loop (Fig. 1). These loops can be either stabilizing or destabilizing for structure formation.

In the Turner 2004 model, internal loop stabilities are modeled using sequence-dependent stabilities that depend on the identity of the first mismatch, loop size, and loop asymmetry^15^. We provide 1×1, 1×2, and 2×2 loops (referring to the number of unpaired nucleotides on each side of the loop) in lookup tables that contain experimental values when available and fitted values when the experiments are not available.

The loop motif stabilities are in Table S14. We found that 1mΨ-1mΨ and 1mΨ -U first mismatches stabilize internal loop formation as compared to U-U first mismatches. A first 1mΨ-1mΨ mismatch stabilizes by an additional −0.92 ± 0.12 kcal/mol, based on seven experiments. The first 1mΨ-U mismatch stabilizes by an additional −1.41 ± 0.15 kcal/mol, based on three experiments. The additional stability does not apply to U-1mΨ first mismatches (where the U is 3’ to the adjacent pair and the 1mΨ is 5’ to the adjacent pair).

We also observe that closing an internal loop with an A-1mΨ or a 1mΨ-A pair destabilizes internal loop formation to a greater extent than an analogous A-U closure. Across 25 experiments, the average destabilization is 0.70 ± 0.10 kcal/mol.

To model internal loop stabilities we use the Turner 2004 model, adding the additional terms for 1mΨ -U or 1mΨ-1mΨ first mismatches and A-1mΨ or 1mΨ-A closure. For 1×1, 1×2, and 2×2 loops, we insert the experimental value when it is available. Using the model with the three additional terms, we estimate the ΔG°_37_ for these internal loops with a root-mean-squared deviation of 0.92 kcal/mol and a mean absolute deviation of 0.68 kcal/mol.

Four additional duplexes, each containing two 1mΨ-C single mismatches (a.k.a. 1×1 internal loops), were studied by optical melting to test this model and to test for non-nearest neighbor effects (Table S15). For these duplexes, the ΔΔG°_37_ between the experiment and the nearest neighbor estimate ranged between −0.85 and 0.16 kcal/mol, a deviation of less than 7.3% of the total helix ΔG°_37_ (Figure 7 and Table S16). This shows excellent agreement with nearest neighbor estimates.

**Figure 7.**
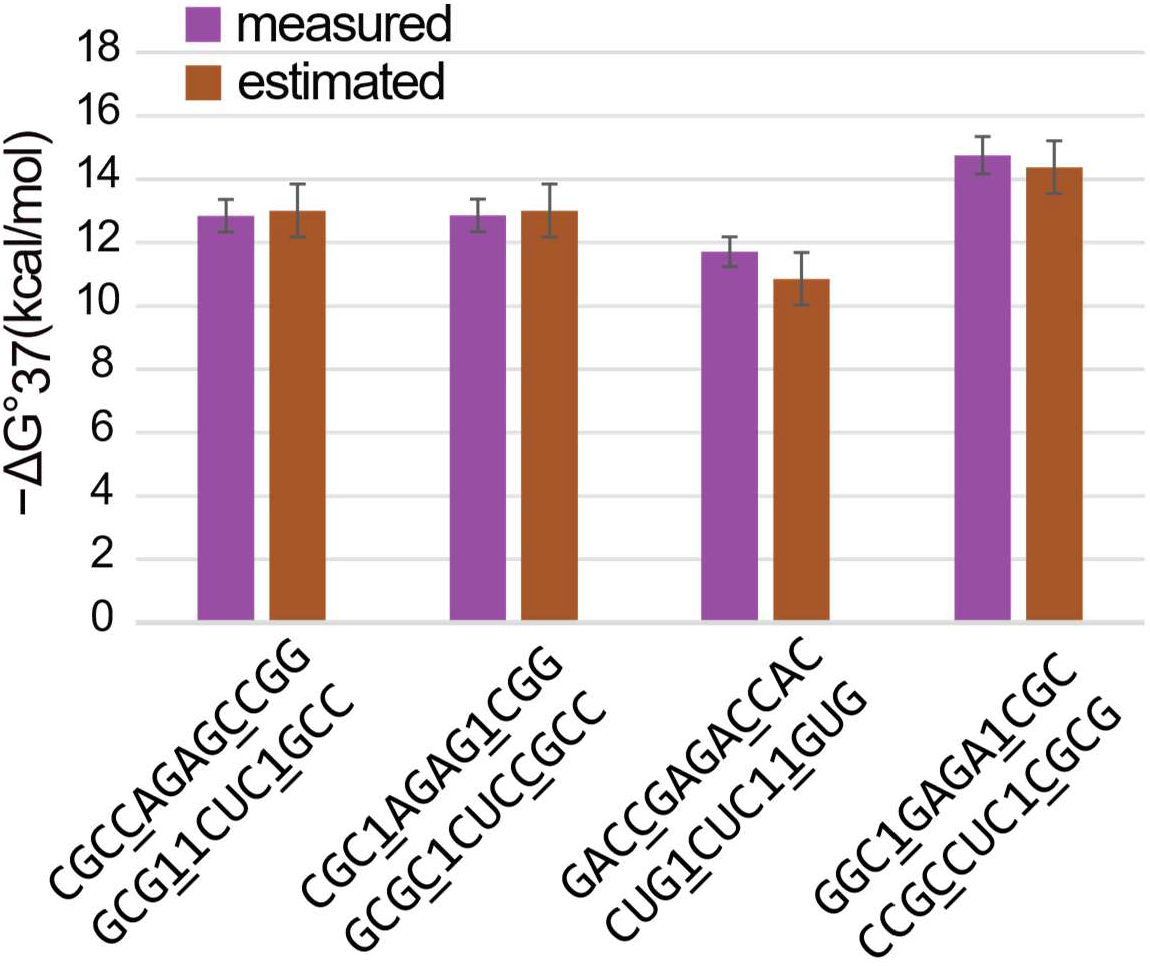
Agreement between measured and estimated ΔG°_37_ for duplexes with two 1×1 internal loops (details in Table S16).

### Bulge loops

The stabilities for four single nucleotide bulge loop model systems were measured by optical melting where 1mΨ was the bulge nucleotide or at least one closing base pair was 1mΨ-A (Table S17). Bulge loops are an interruption of the helix with one or more unpaired nucleotides on one side of the loop (Fig. 1). The Turner 2004 parameters treat these as sequence independent, with a stability penalty that grows with bulge loop size^15^. For single nucleotide bulges, the model also adds the helical stack of the adjacent pairs, which assumes that the single bulge is flipped out of the helix.

For three of the systems, the standard Turner 2004 model was able to estimate the ΔG°_37_ (Table S18); the differences between the estimates and the experiments are −0.17±0.85, 0.24±0.94, and 0.75±0.77 kcal/mol. We therefore use the model unchanged for 1mΨ. For one duplex containing a bulged 1mΨ flanked by two GC pairs, however, the model underestimates the stability by 2.55±0.80 kcal/mol. It is unclear why 1mΨ stabilizes this duplex.

### Modeling of tRNA Secondary Structures Ensembles is Improved

To test the 1mΨ parameter set, we predicted the folding ensembles of tRNAs using 1mΨ parameters or using U parameters for the 1mΨ nucleotides. We previously demonstrated that the normalized ensemble defect (NED), which is the average probability that nucleotides do not fold to the correct structure, characterizes the tRNA folding ensemble and distinguishes between sequences that will correctly fold to the cloverleaf from those that will not^34, 35^. NED ranges from 0, a perfect cloverleaf, to 1, a sequence that has zero probability of forming the correct pairs and loops^36^. NED therefore provides a benchmark for quantifying the improvement of modeling tRNA structure with 1mΨ as compared to U parameters. Additionally, we can estimate structures from pairing probabilities using maximum expected accuracy (MEA) structure prediction, which assembles structures from the most probable pairs^16, 37^.

For the 55 tRNA sequences from the Sprinzl database that contain 1mΨ, we find that the NED significantly improves from 0.267 to 0.252, P=3.0×10^-6^, by treating the 1mΨ correctly using our new parameters^38^. For predicted MEA structures, the sensitivity (percent of known pairs correctly predicted) significantly improves from 86.0 to 87.5% (P=0.026). The positive predictive value (percent of predicted pairs in the known structure) improves from 81.4 to 82.5%, but the change is not significant (P=0.081). Fig. 8 shows an example where the 1mΨ parameters dramatically improve the structure prediction accuracy. Table S19 provides the results across all sequences.

**Figure 8.**
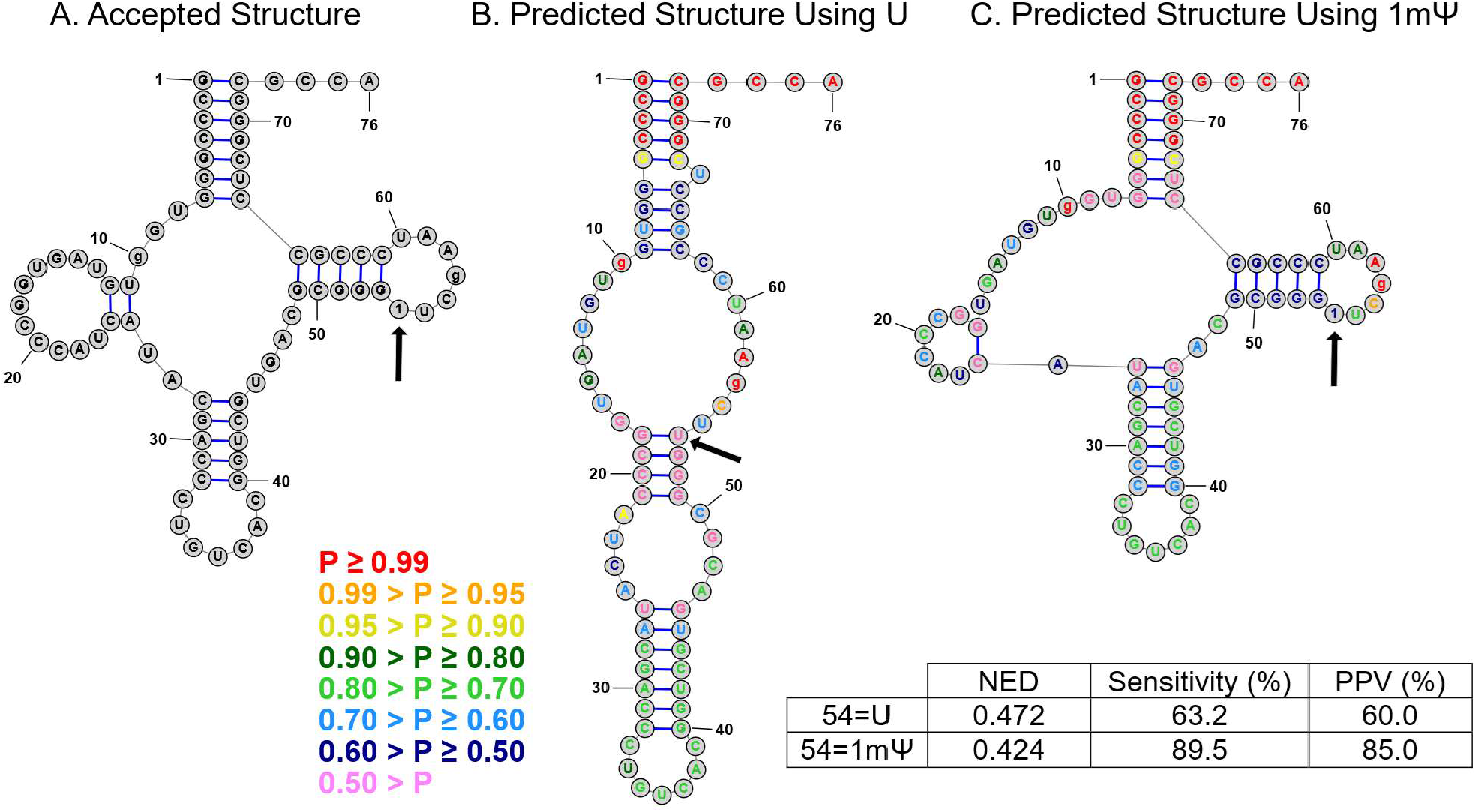
1mΨ parameters improve the accuracy of the folding ensemble for the *Haloferax volcanii* Asp CUC tRNA. Panel A is the structure from the Sprinzl database ^38^. Panel B shows the MEA structure prediction using U for position 54, which is known to be 1mΨ. Panel C shows the MEA structure prediction where position 54 is correctly treated as 1mΨ. Predicted base pairing probabilities are annotated by color as shown in the legend. The normalized ensemble defect improves from 0.472 to 0.424. The sensitivity (percent of known pairs correctly predicted) improves from 63.2 to 89.5%. The positive predictive value improves from 60.0 to 85.0%. In the predictions, the N2,N2-dimethylguanosine at position 10 and the 1-methylinosine at position 57 are treated as G, but not allowed to base pair. These figures were prepared with the RNAstructure structure editor.

## Discussion

### Parameter Availability

The nearest neighbor parameters are freely provided as part of the RNAstructure software package (https://rna.urmc.rochester.edu/RNAstructure.html) in plain text files^28^. Most command-line programs can be invoked in RNAstructure using the --alphabet parameter, which can specify alternative alphabets to the canonical RNA or DNA nucleotides, including m6A, DNA P and Z, or 1mΨ. The 1mΨ parameters are available in two alphabets. The first is A, C, G, U, and 1mΨ (invoked with --alphabet 1). The second is A, C, G, and 1mΨ (invoked with --alphabet Full1), which is designed for use with sequences that are fully substituted with 1mΨ for U. This is of special interest to the community because mRNA vaccines are fully substituted with 1mΨ^39^. The Full1 alphabet conveniently accepts U in the sequence for 1mΨ and is faster for calculations because the number of the nearest neighbor terms is smaller.

### 1mΨ Substitution Generally Stabilizes Folding

As compared to U, 1mΨ substitution generally stabilizes folding stability. For helical stacks, some notable exceptions occur for A-1mΨ pairs adjacent to G-U pairs (Figure 4 and Table S4). The changes in stabilities are sequence-dependent, requiring the nearest neighbor approach for estimating stabilities (Fig. 3 and Table S3). Figure 9 demonstrates the use of the parameters and shows how stabilizing substitutions are for a helix.

**Figure 9.**
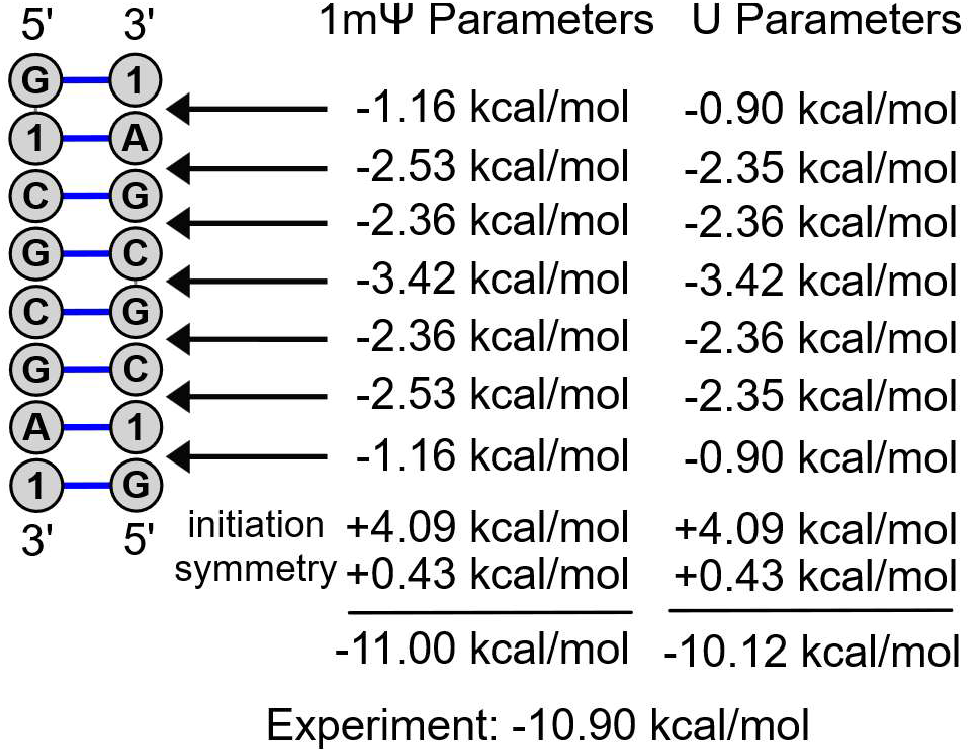
Example calculation for a self-complementary helix^11^. The left calculation uses the 1mΨ parameters and the right calculation estimates the stability with U^13, 30^. The helix has seven base pair stacks. The first contains a G-1mΨ pair followed by a 1mΨ-A pair. Its stability is available in Table S4. The second stack is the 1mΨ-A pair followed by a C-G pair, and its stability is in Table S3. In addition to the helix stacks, an intermolecular initiation penalty and a symmetry penalty are applied. The experimentally measured stability (Table S5) is −10.90 kcal/mol and is better estimated using the 1mΨ parameters.

Beyond stabilizing base pairs, substituting 1mΨ for U in loops is also generally stabilizing when the 1mΨ is able to stack on adjacent base pairs. In internal loops, we find empirically that 1mΨ-U and 1mΨ-1mΨ first mismatches provide additional stability of −1.41±0.15 and −0.93±0.12 kcal/mol, respectively, compared to an analogous U-U mismatch. For hairpin, multibranch, and internal loops, terminal mismatches and dangling ends involving 1mΨ are also generally more stabilizing than those with an analogous U. Again, 1mΨ-U tends to stabilize; in the example of a 1mΨ-U following a G-C pair, the additional stability is −1.10±0.66 kcal/mol. The greater folding stability of 1mΨ, which is present in many mRNA vaccines, is consistent with previous reports that more stable RNAs make better mRNA vaccines^26, 40, 41^.

### Comparison to Previous Parameters

Moderna had previously reported a set of the seven stacking nearest neighbor parameters for fully substituted sequences, using experiments performed by DNA Software^26^. Our work expands on this by considering a mixture of U and 1mΨ in sequences and by also including 1mΨ-G base pairs. Figure 10 and Table S20 compare the two sets of parameters, which agree within 0.4 kcal for each parameter, supporting the accuracy of the parameters provided herein.

**Figure 10.**
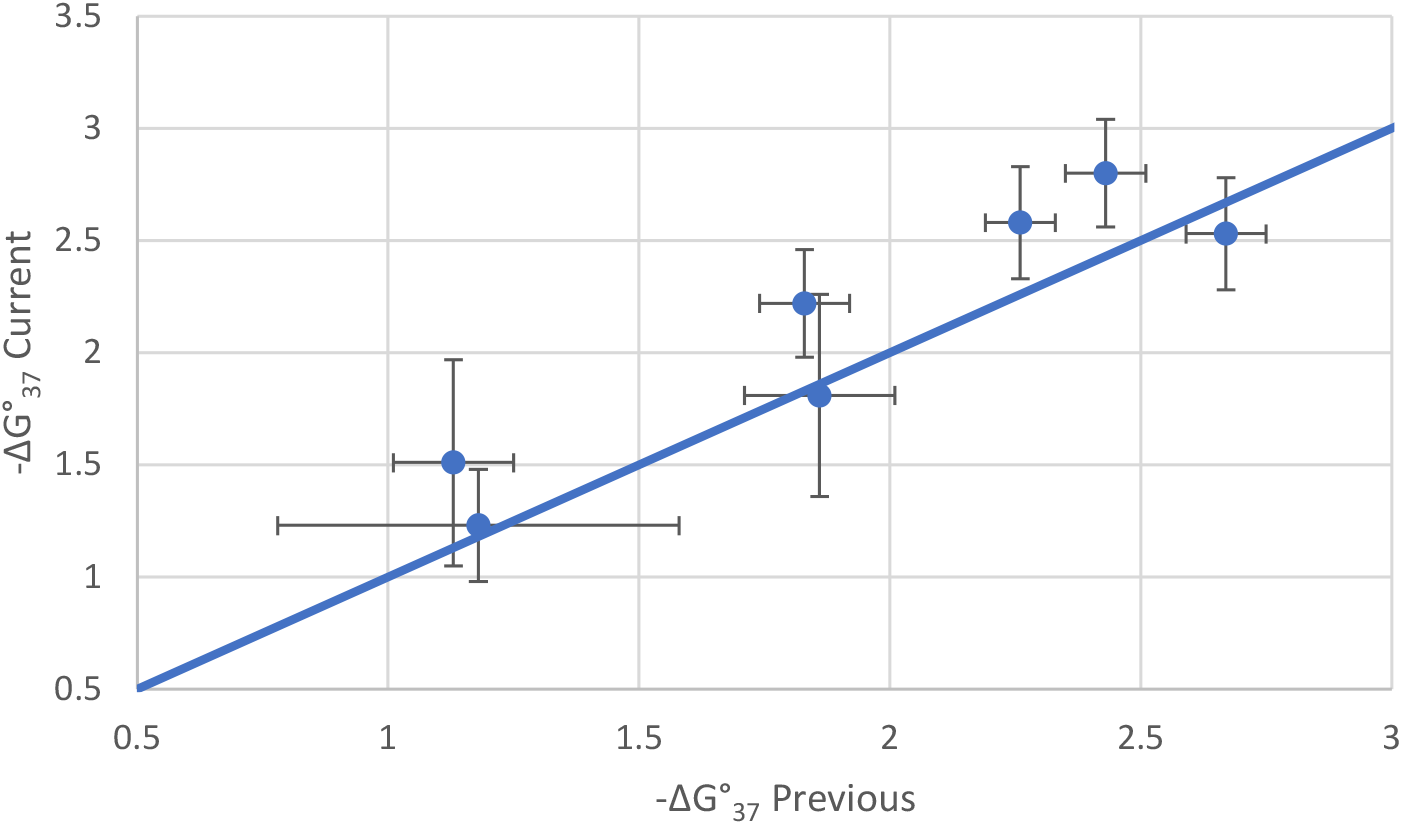
Agreement for fully substituted stacks between prior parameters^26^ and these parameters. The diagonal line is shown.

### Reproducibility of Optical Melting

As part of this study, we performed optical melting at three sites to compare results and to prior literature values from the University of Rochester. Overall, the agreement was excellent with an average of 2.1% uncertainty in ΔG°_37_. We recommend that future studies use the set of duplexes here (Table S1) to test reproducibility and ensure that free energy changes are comparable across sites and across time to previous publications.

### Enthalpy Changes

In this work, we focused on determining parameters to estimate ΔG°_37_, the folding free energy change at 37 °C. For the canonical RNA alphabet, a set of enthalpy change (ΔH°) nearest neighbor parameters are available that can be used to extrapolate folding free energy changes to temperatures between 10 and 60 °C^42^. Future work could derive a set of ΔH° parameters that include 1mΨ. The uncertainties for ΔH°, however, are larger than ΔG°_37_. It was previously noted that the reproducibility for optical melting estimates for DNA duplexes across sites is 3%, 6%, and 6% for ΔG°_37_, enthalpy change, and entropy change, respectively^13, 29^. For the three RNA calibration duplexes studied here, we found average uncertainties of 2.12%, 14.1%, and 16.3% in enthalpy changes (Table S1). In part, the smaller uncertainties for ΔG°_37_ as compared to the larger uncertainties for enthalpy and entropy changes result from the large (0.999) correlation of enthalpy and entropy changes^13^. Because of the larger uncertainties in enthalpy changes, additional optical melting experiments would likely be needed to fit enthalpy change parameters.

### Applications

We demonstrated using tRNAs that these nearest neighbor parameters for 1mΨ improve the modeling of RNA sequences where 1mΨ is known to naturally occur. Another important application of these parameters is the design of mRNA sequences. 1mΨ is commonly used to substitute for U in mRNA vaccines and therapies to evade the innate immune response^7^. Additionally, we know that mRNA open reading frames (ORFs) designed to both optimize codon usage and to maximize base pairing (by minimizing the folding free energy change) can increase protein expression^26, 41, 43, 44^. An important factor in this increased expression is the stabilization of the RNA against spontaneous hydrolysis by the inline geometry that is more easily sampled in loops^40^, and 1mΨ substitution is known to increase mRNA half life^45^. These parameters will improve modeling of structures of mRNA designs as compared to estimates relying on folding thermodynamics of U. An additional application of these parameters is the design of sequences for RNA nanostructures^24, 46, 47^. Selective incorporation of 1mΨ using a five-nucleotide alphabet could be used to optimize the folding of structures against alternative structures. For example, the selective use of 1mΨ-1mΨ for the first mismatches in the loops of sequence designs would stabilize the target structure relative to U-U interactions.

## Methods

### Preparation of model RNA duplexes at the Institute of Bioorganic Chemistry

Oligonucleotides were synthesized on a BioAutomation MerMade12 DNA/RNA synthesizer using β-cyanoethyl phosphoramidite chemistry and commercially available phosphoramidites (ChemGenes, GenePharma, or Glen Research), where standard protocols were followed as described previously^48, 49^. For deprotection, oligoribonucleotides were treated with a mixture of 30% aqueous ammonia and ethanol (3:1 v/v) for 16 h at 55°C. Silyl protecting groups were removed with the use of triethylamine trihydrofluoride. The deprotected oligonucleotides were purified using silica gel TLC in a mixture of 1-propanol, aqueous ammonia, and water (55:35:10 v/v/v), as described previously^50^. Mass spectrometry analyses (MALDI) was performed for most of the oligonucleotides.

### UV melting experiments at the Institute of Bioorganic Chemistry

The thermodynamic measurements were performed for nine various concentrations in the range 100 µM - 1 µM on a JASCO V-650 UV/Vis spectrophotometer in buffer containing 1 M sodium chloride, 20 mM sodium cacodylate, and 0.5 mM Na_2_EDTA, pH 7^13, 51^. Oligonucleotide single-strand concentrations were calculated from the absorbance above 80 °C, and single strand extinction coefficients were approximated by a nearest-neighbor model^52, 53^. Absorbance vs. temperature melting curves were measured at 260 nm with a heating rate of 1 °C/min from 0 to 90 °C on a JASCO V-650 spectrophotometer with a thermoprogrammer. The melting curves were analyzed, and the thermodynamic parameters were calculated from a two-state model with the program MeltWin 3.5^54^. For most duplexes, the ΔHº derived from T_M_^-1^ vs. ln(C_T_/4) plots are within 15% of that derived from averaging the fits to individual melting curves, as expected if the two-state model is reasonable.

### Temperature calibration of instruments at the Institute of Bioorganic Chemistry

Temperature calibration was performed on the JASCO V-650 UV/Vis spectrophotometer using a buffer containing 1 M sodium chloride, 20 mM sodium cacodylate, and 0.5 mM Na_2_EDTA, pH 7, which was used for thermodynamic measurements. To acquire the correct temperature, a needle thermocouple was placed into 300 μl (1cm pathlength) quartz cuvette and covered with thermoisolating fabrics. The thermocouple was previously calibrated by placing it in wet ice and boiling water. The temperature calibration was acquired while running the program used for standard thermodynamic measurements and performed between 4 and 90 °C at a heating rate of 1 °C /min. The temperature indicated by thermocouple and instrument was recorded on two of six positions (positions two and five) of the JASCO cuvettes holder. Those measurements were performed three times for each of the two positions.

Optical melting curves were then corrected for the temperature deviation with post-processing. A C++ program was written to linearly extrapolate the temperature recorded by the instrument to that determined by thermocouple, using the adjacent two temperature readings. This corrected temperature was used in MeltWin for the data analysis.

### UV melting experiments at DNA Software, Inc

The thermodynamic measurements were performed on a Beckman DU800 UV/Vis spectrophotometer with thermoprogrammer. For each set of sequences (i.e., duplex or single strands), six strand concentrations in the range 100 µM - 1 µM were measured in a buffer containing 1 M sodium chloride, 20 mM sodium cacodylate, and 0.5 mM Na_2_EDTA, pH 7. Oligonucleotide single-strand concentrations were calculated from the absorbance above 80 °C, and single-strand extinction coefficients were approximated by a nearest-neighbor model. Absorbance vs. temperature melting curves were measured at 260 nm with a heating rate of 1 °C/min from 5 to 90 °C. The melting curves were analyzed, and the thermodynamic parameters were calculated from a two-state model with the program MeltWin 3.5. For most duplexes, the ΔHº derived from T_M_^-1^ vs. ln(C_T_/4) plots are within 15% of that derived from averaging the fits to individual melting curves, as expected if the two-state model is reasonable.

### Temperature calibration of instruments at DNA Software, Inc

Temperature calibration was performed on a Beckman DU-800 UV/Vis spectrophotometer. To acquire the correct temperature, a type K thermocouple was placed into 300 μl (1 cm pathlength) quartz cuvette filled with 250 μl distilled water and sealed with parafilm to minimize evaporation. The temperature calibration was acquired while running the program used for standard thermodynamic measurements and performed between 5 and 90 °C at a heating rate of 1 °C /min. The temperature indicated by thermocouple and instrument was recorded on two of six positions (positions two and five) of the Beckman DU800 cuvettes holder. Those measurements were performed twice for each of the two positions.

Optical melting curves were then corrected for the temperature deviation with post-processing. The experimental linear correction equation was determined by linear regression:

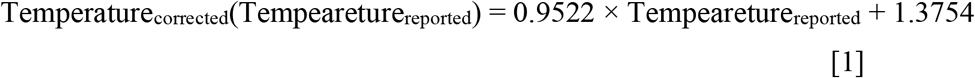

where Temperature_corrected_ is the revised temperature used for fitting the thermodynamics and Tempeareture_reported_ is the temperature reported by the instrument.

After applying this temperature correction, the Temperature_corrected_ agrees with the actual thermocouple temperature with a slope of 1.000, intercept of 0.000 °C, and an average deviation of 0.1938 °C. The correction equation was applied to the experimental UV melting data using Excel. Temperature_corrected_ was used in MeltWin for the data analysis.

### Propagation of uncertainties

Nearest neighbor parameter values are extrapolated from linear equations:

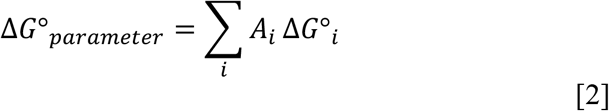

where A_i_ are coefficients and ΔG°_i_ are nearest neighbor parameters or experimental results. To propagate uncertainties, we use the standard approach, assuming uncertainties in parameters are uncorrelated^55^:

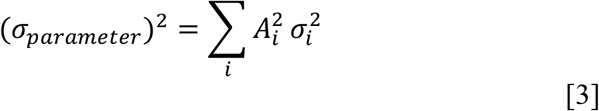

where σ_i_ is the uncertainty of ΔG°_i_. For optical melting experiments:

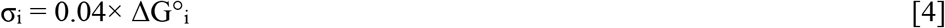

following Xia et al.^13^. This uncertainty was chosen as a conservative estimate based on the observation that DNA optical melting experiments were consistent to within 3% across sites^29^.

### Helical stack parameters

For duplexes to determine helical parameters, we generally chose sequences that had been previously studied with U and had optical melting curves that were consistent with two state denaturation^13, 30^. Helical stack parameters, including all base pair stacks with A-1mΨ or G-1mΨ pairs, were fit by linear regression. Table S2 shows the set of helices studied by optical melting. The canonical pair stacks were fixed at the values of Xia et al. for Watson-Crick-Franklin pairs and Chen et al. for G-U base pairs^13, 30^. The values of the canonical pair stacks were subtracted from the measured helical stability, leaving the stability attributed to the stacks incorporating 1mΨ. These values were then fit with non-error-weighted linear regression using the Statsmodels library in a custom Python program (Tables S3 and S4)^56^. Uncertainty estimates are the standard error of the regression values. The R^2^ for the fit was 0.995.

We determined that no terminal stability penalties were needed for 1mΨ-A and 1mΨ-G pairs. When those terms were included in the linear regression, the terminal 1mΨ-A term was - 0.12 ± 0.21 kcal/mol and the terminal 1mΨ-G penalty was 0.01 ± 0.19 kcal/mol. The magnitudes of the fitted penalties are small, and the uncertainties are larger than the value.

Tables S3 and S4 provide the coverage of individual stacks in the fitting dataset. For stacks with 1mΨ-A pairs, the coverage is excellent with a maximum of 25 occurrences for A-1mΨ followed by G-C and a minimum of 5 occurrences for 1mΨ-A followed by 1mΨ-A. Because of the large number of helical stacking parameters for G-1mΨ pairs, the coverage is lower with a maximum of 14 occurrences of 1mΨ-G followed by G-C and a minimum of 2 for both 1mΨ-G followed by C-G and for 1mΨ-A followed by G-U.

### Dangling end parameters

Dangling ends, i.e. an unpaired nucleotide at the end of a helix, is known to stabilize helix formation. A set of 20 duplexes were studied to determine dangling end parameter values (Table S7). Two duplexes had a difference in ΔH° between the two fit methods is greater than 15%, which can indicate non-two-state melting. Because only 20 dangling ends were melted of 52 possibilities involving 1mΨ, we decided to include the data from these melts for subsequent analysis.

From the optical melting experiments, the dangling end stabilities are determined using:

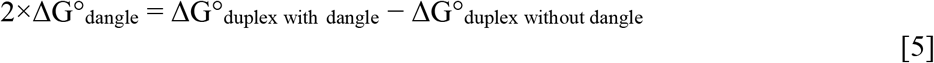

where ΔG°_dangle_ is a parameter that depends on the sequence of the terminal base pair, the sequence of the dangling end, and the position of the dangling end (as 5’ or 3’ to the paired strand). These results are available in Table S9. The duplexes in this study are self-complementary and, therefore, contain the dangle at each end of the helix, which accounts for the factor of two on the left of the equation. ΔG°_duplex without dangle_ is the helix without the dangling end. When available, an experimental value is used. When it is not available, this stability is estimated with the helical stack parameters.

Table S9 shows the dangling end parameter values. When available, the nearest neighbor parameters use the experimental result for the dangling ends. The 3’ dangling G on a terminal 1mΨ-A pair was measured twice, and the mean value is used. Two duplexes had dangling ends that were destabilizing by equation 5, AUGCA(1mΨ)(1mΨ)_2_ and C(1mΨ)CAUGA_2_. This is unexpected, and both have a ΔG°_duplex without dangle_ estimated using nearest neighbor parameters. Therefore, we excluded these sequences from the analysis.

For dangling ends that were not studied, we extrapolated values. First, we calculated:

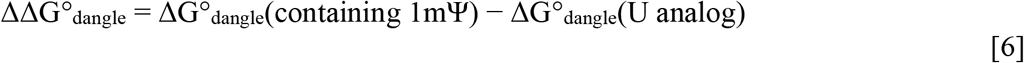

where ΔG°_dangle_(containing 1mΨ) is the value of the dangle as estimated from an experiment using equation 5 and ΔG°_dangle_(U analog) is the value of the dangle for the analogous dangling end that has U replacing 1mΨ as found in the Turner 2004 rules and tabulated by Zuber et al.^22^.

The mean ΔΔG°_dangle_ for measured 1mΨ 5’ dangling ends on canonical terminal pairs was −0.22 ± 0.07 kcal/mol. Similarly, the mean ΔΔG°_dangle_ for measured 1mΨ 3’ dangling ends on canonical terminal pairs was +0.10 ± 0.07 kcal/mol. To extrapolate dangling 1mΨ on canonical base pairs, we added the ΔΔG°_dangle_ (5’ or 3’ as applicable) to the analogous U dangling end.

To extrapolate the term for a 5’ dangling U on an A-1mΨ pair, we added the mean ΔΔG°_dangle_ for 5’ dangling A, C, and G on an A-1mΨ pair of −0.16 ± 0.02 kcal/mol to the stability of a 5’ dangling U on an A-U pair. To extrapolate the terms for 3’ dangling C, G, and U on an A-1mΨ pair, we added the ΔΔG°_dangle_ for the 3’ dangling A, 0.49 ± 0.46 kcal/mol, to the analogous dangle on an A-U pair.

To extrapolate the terms for 5’ dangling A, C, G, and U on an 1mΨ-A pair, we added the mean ΔΔG°_dangle_ for 5’ dangling A, C, and G on an A-1mΨ pair of −0.16 ± 0.02 kcal/mol to the analogous dangle on a U-A pair. To extrapolate the stability of a 3’ dangling U on a 1mΨ-A pair, we added the mean ΔΔG°_dangle_ for 3’ dangling A, C, and G on 1mΨ-A of −0.43 ± 0.02 kcal/mol to the 3’ dangling U on a U-A pair.

In the Turner 2004 model, dangling ends on G-U and U-G pairs are set equal to the equivalent dangle on an A-U or U-A pair, respectively. We use this approach for 1mΨ dangling ends on terminal G-U and U-G pairs. We also use dangling ends for terminal A-1mΨ and 1mΨ-A to model the dangling ends on terminal G-1mΨ and 1mΨ-G pairs.

### Terminal mismatch parameters

Terminal mismatches, a non-canonical pair adjacent to the end of a helix is also known to be stabilizing. These terms are sequence dependent for the closing base pair and the mismatch. Using a set of 40 duplexes, we modeled the stability of all the terminal mismatches that contain 1mΨ.

From the experiments, the terminal mismatch stability is determined by:

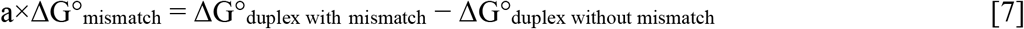

where ΔG°_duplex with mismatch_ is the stability determined by optical melting for the duplex that has the mismatch. ΔG°_duplex without mismatch_ is the stability of the core helix without the terminal mismatch. When available, we used the experimentally determined value. We used nearest neighbor estimates when it was not studied experimentally. Most duplexes in this study were composed of self-complementary strands and, therefore, contain the terminal mismatch at each end of the duplex. For these, a = 2. For three experiments, the duplex strands were non-self-complementary. For these, a = 1 because the duplex has a single terminal mismatch.

Table S10 provides the terminal mismatch parameter values. When available, we use the experimental value. For a subset of terminal mismatch sequences, multiple experiments were performed, and we use the mean value.

To extrapolate parameters that were not measured, we compared the stability of mismatches containing 1mΨ to analogous mismatches with U:

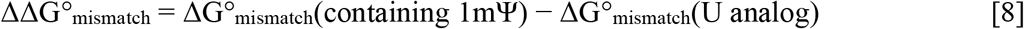

where ΔΔG°_mismatch_ is the difference in stability between the mismatch with 1mΨ and a mismatch with U in the analogous position, using Turner 2004 rules as tabulated by Zuber et al. (2017).

For a C-1mΨ following an A-U or U-A pair, we added the mean ΔΔG°_mismatch_ for C-1mΨ following C-G and G-C pairs (−0.23 ± 0.28 kcal/mol) to the analogous C-U mismatch stability. For a 1mΨ-1mΨ mismatch following an A-U or U-A pair and for an U-1mΨ mismatch following A-U, C-G, G-C, or U-A pairs, we added the mean ΔΔG°_mismatch_ for a 1mΨ-1mΨ mismatch following C-G and G-C pairs (−0.33 ± 0.33 kcal/mol) to the stability of the analogous U-U mismatch. For a 1mΨ-C mismatch following an A-U pair or a U-A pair, we added the mean ΔΔG°_mismatch_ for a 1mΨ-C mismatch following a C-G or G-C pair (−0.60 ± 0.46 kcal/mol) to the analogous U-C mismatch. For a 1mΨ-U mismatch following an A-U pair, we added the ΔΔG°_mismatch_ for a 1mΨ-U following a U-A pair to the analogous U-U mismatch. For a 1mΨ-U mismatch following a C-G pair, we added the ΔΔG°_mismatch_ for a G-C pair to the analogous U-U mismatch.

To extrapolate the stability of a C-U mismatch following an A-1mΨ pair, we subtracted the ΔΔG°_mismatch_ for a C-1mΨ mismatch following an A-U pair from the stability of a C-1mΨ mismatch following a A-1mΨ pair. The stability of a U-C following an A-1mΨ pair was estimated by subtracting the mean stability of a 1mΨ-C mismatch on C-G and G-C closing pairs (−0.60 ± 0.46 kcal/mol) from the stability of a 1mΨ-C mismatch on an A-1mΨ pair. Likewise, the stability of a U-U mismatch following an A-1mΨ pair was estimated by subtracting the mean stability of a 1mΨ-1mΨ mismatch on C-G and G-C closing pairs (−0.33 ± 0.33 kcal/mol) from the stability of a 1mΨ-1mΨ mismatch on an A-1mΨ pair. The stability of a 1mΨ-U mismatch following an A-1mΨ pair is extrapolated as the sum of the ΔΔG°_mismatch_ for a 1mΨ-U mismatch following an A-U pair plus the stability of U-U mismatch following an A-1mΨ pair.

To extrapolate the stability of a U-1mΨ mismatch following an A-1mΨ pair, we summed the dangling end stabilities of a 3’ dangling U and the 5’ dangling 1mΨ on an A-1mΨ pair. This follows an extrapolation used for purine-pyrimidine terminal mismatches in the Turner 2004 rules^57^. Similarly, this sum of dangling ends approach was used for a U-1mΨ mismatch following a 1mΨ-A pair.

The stability of a C-U mismatch following a 1mΨ-A pair is estimated by subtracting the ΔΔG°_mismatch_ for a C-1mΨ mismatch following an A-U pair from the stability of a C-1mΨ mismatch following a 1mΨ-A pair. Similarly, the stability of a U-C mismatch following a 1mΨ-A pair is estimated by subtracting the ΔΔG°_mismatch_ for a 1mΨ-C mismatch following an A-U pair from the stability of a 1mΨ-C mismatch following a 1mΨ-A pair. The stability of a 1mΨ -U mismatch following a 1mΨ-A pair is estimated as the ΔΔG°_mismatch_ for a 1mΨ -U mismatch following a U-A pair plus the stability of a U-U mismatch following a 1mΨ-A pair.

In the Turner 2004 rules, for mismatches following G-U or U-G pairs, when not available experimentally, we use the stability for the analogous mismatch on an A-U or U-A pair, respectively. We used this approach here for mismatches on G-U or U-G pairs. Similarly, we used mismatches on 1mΨ-A or A-1mΨ pairs to estimate mismatch stabilities on 1mΨ-G or G-1mΨ pairs, respectively.

### Hairpin loops

Hairpin loop stabilities are determined from hairpin stem-loop experiments using:

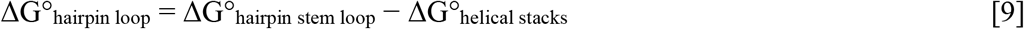

where ΔG°_hairpin stem loop_ is the experimentally determined stability of the hairpin with its closing stem, and ΔG°_helical stacks_ is the total stability of the helix stacking terms including terminal pair penalties but excluding intermolecular initiation.

The set of experimentally studied hairpin stem-loop structures is contained in Table S11, and the hairpin loop stabilities are in Table S12. The hairpin loop nearest neighbor model is:

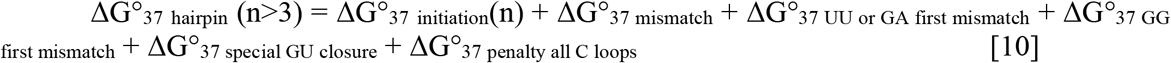

where n is the number of unpaired nucleotides in the loop. ΔG°_37 initiation_(n) is a sequence-independent term that is the cost for closing the loop of n nucleotides and is a lookup based on fit to experiments. ΔG°_37 mismatch_ is the terminal mismatch term for the first mismatch adjacent to the closing base pair. ΔG°_37 UU or GA first mismatch_ and ΔG°_37 GG first mismatch_ are empirical bonuses for UU, GA, or GG first mismatches that were observed in optical melting experiments. Hairpin loops closed by G-U pairs with two preceding G residues receive a bonus, ΔG°_37 special GU closure_. Loops composed entirely of C residues receive a penalty, ΔG°_37 penalty all C loops_. We determined that ΔG°_37 UU or GA first mismatch_ applies to loops where the first mismatch is 1mΨ-1mΨ, 1mΨ-U, or U-1mΨ.

### Internal loops and bulge loops

Internal loop and bulge loop stabilities are determined from experiments using one of two equations. When the helix without the loop has been measured, the loop stability is determined with:

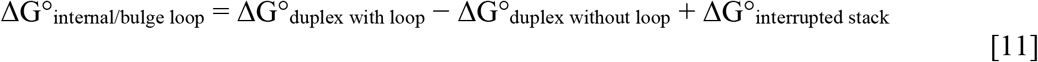

where ΔG°_duplex with loop_ and ΔG°_duplex without loop_ are the stabilities determined by optical melting experiments^58^. ΔG°_interrupted stack_ is the stability of the helical stacking parameter for the stack between adjacent pairs in the duplex without the loop that is interrupted by the loop. The A-U terminal pair penalty is additionally applied to ΔG°_interrupted stack_ per A-U pair in the stack. ΔG°_interrupted stack_ accounts for the fact that the internal loop divides the helix found without the loop into two separate helices.

When the helix without the loop has not been measured, the loop stability is determined with:

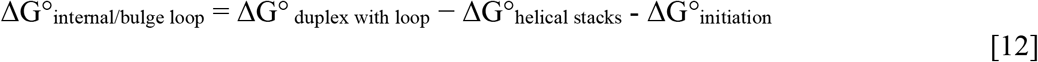

where ΔG° _duplex with loop_ is measured by optical melting experiments^59^. ΔG°_helical stacks_ is the sum of the helical stacking parameters in the two helices plus A-U terminal pair penalties for each A-U helix end (adjacent to the internal loop or at the end of the duplex). ΔG°_initiation_ is the intermolecular initiation term.

Tables S13 and S15 provide the set of internal loops studied by optical melting. Table S14 provides the set of internal loop stabilities used to inform the model. Table S16 provides a set of duplex stabilities that tested the model.

### Multibranch and exterior loops

Multibranch and exterior loop stabilities are estimated with sequence-independent terms that are unchanged, dangling end terms, mismatch terms, and coaxial stacking terms^11, 60^. The stability for coaxial stacking for directly adjacent helices is the stability of a helix stack parameter^61^. The stability for coaxial stacking of helices separated by one mismatch is the sum of a sequence-independent term where the backbone is not continuous and the terminal mismatch parameter where the helix is continuous^14, 62^.

### tRNA structure predictions

We used the Sprinzl database of tRNA sequences and structures and extracted the 55 entries with 1mΨ^38^. For modifications other than 1mΨ, we mapped nucleotides back to their unmodified base^4, 14^. For modifications to the Watson-Crick face of the base or that make the base ring non-planar, we prevent the base from forming base pairs in structure prediction. The modifications for which we forbid pairing are: N2,N2-dimethylguanosine, 1-methylinosine, dihydrouridine, galactosyl-queuosine, queuosine, archaeosine, 3-(3-amino-3-carboxypropyl)uridine, and 1-methylguanosine.

We predicted the secondary structures and base pairing probabilities with RNAstructure using the 1mΨ alphabet of nearest neighbor parameters (for both U and 1mΨ version of the sequence). We tested significance using a paired t-test with the ttest_rel function from SciPy^63^. We used a 1-tailed test (to test the null hypothesis that the 1mΨ parameters do not improve modeling performance compared to U parameters) with α = 0.05.

## Acknowledgements

This work was partially supported by National Institutes of Health grants R35GM145283 to D.H.M., R21GM148835 to S.A., R35GM127064 to P.C.B., and R15GM085699 to B.M.Z., by a contract from Moderna to E.K. and R.K, and a contract by Moderna to J.S. Research was also supported by grants 2022/45/B/ST4/03586 (to R.K.) and 2021/41/B/NZ1/03819 and 2020/39/B/NZ1/03054 (to E.K.) from the National Science Center of Poland. O.H. was partially supported by National Institutes of Health training grant T32GM135134.

## Notes

https://rna.urmc.rochester.edu/RNAstructure.html

### Competing Interest Statement

Haining Lin and Mihir Metkar are employees of and shareholders in Moderna, Inc.
John SantaLucia and Holly SantaLucia are employees of and shareholders in DNA Software, Inc.
David Mathews has an equity stake in Coderna.AI.

