## Supplemental Figures and Tables for "RNA Folding Nearest Neighbor Parameters Including the Modification 1-Methyl-Pseudouridine"

Table S1. Agreement between calibration duplex optical melting data. The buffer used was 1 M sodium chloride, 20 mM sodium cacodylate, 0.5 mM $Na_2EDTA$, pH 7.

| Site: | Duplex | **Average of curve fits** | | | | **$T_M^{-1}$ vs log $C_T$ plots** | | | |
|---|---|---|---|---|---|---|---|---|---|
| | | **$-\Delta H^\circ$ (kcal/mol)** | **$-\Delta S^\circ$ (eu)** | **$-\Delta G^\circ_{37}$ (kcal/mol)** | **$T_M$[a] (°C)** | **$-\Delta H^\circ$ (kcal/mol)** | **$-\Delta S^\circ$ (eu)** | **$-\Delta G^\circ_{37}$ (kcal/mol)** | **$T_M$[a] (°C)** |
| Poznan | CGCGCG<br>GCGCGC | 58.4±5.3 | 157.5±16.2 | 9.58±0.33 | 59.0 | 56.0±2.8 | 150.3±10.8 | 9.42±0.16 | 59.0 |
| DNA Software | CGCGCG<br>GCGCGC | 52.5±4.0 | 139.4±12.1 | 9.28±0.31 | 59.8 | 41.0±3.8 | 103.9±11.8 | 8.79±0.20 | 62.4 |
| St. Louis | CGCGCG<br>GCGCGC | 60.2±3.6 | 164.2±11.2 | 9.29±0.15 | 56.7 | 62.9±3.0 | 172.6±9.1 | 9.39±0.16 | 56.3 |
| Poznan (2022)[20] | CGCGCG<br>GCGCGC | | | | | 47.8±1.9 | 124.0±5.9 | 9.38±0.14 | 62.8 |
| Rochester (1985)[65] | CGCGCG<br>GCGCGC | | | | | 54.5 | 146.4 | 9.12 | 57.8 |
| % Uncertainty[b] | | | | | | 15.9% | 18.9% | 3.19% | |
| Poznan | GCUACG<br>CGAUGC | 57.8±3.0 | 161.5±9.5 | 7.74±0.17 | 43.5 | 56.4±3.7 | 157.2±11.5 | 7.68±0.11 | 43.2 |
| DNA Software | GCUACG<br>CGAUGC | 63.7±3.1 | 180.5±10.6 | 7.71±0.16 | 42.9 | 79.3±7.3 | 231.2±23.9 | 7.56±0.13 | 41.2 |
| St. Louis | GCUACG<br>CGAUGC | 63.1±8.8 | 179.4±27.9 | 7.69±0.12 | 41.6 | 58.9±5.3 | 166.3±17.0 | 7.36±0.13 | 41.2 |
| Rochester (1998)[13] | GCUACG<br>CGAUGC | | | | | 58.0 | 162.7 | 7.56 | 42.5 |
| % Uncertainty[b] | | | | | | 17.1% | 19.4% | 1.76% | |
| Poznan | GCGUUCGC<br>CGCUUGCG | 68.8±6.3 | 196.3±19.4 | 7.98±0.27 | 47.4 | 65.5±3.0 | 186.0±9.3 | 7.84±0.08 | 47.5 |
| DNA Software | GCGUUCGC<br>CGCUUGCG | 71.1±3.8 | 204.6±12.5 | 7.65±0.11 | 45.8 | 58.9±3.7 | 165.4±12.0 | 7.60±0.06 | 47.5 |
| St. Louis | GCGUUCGC<br>CGCUUGCG | 75.6±5.1 | 218.8±15.5 | 7.69±0.12 | 45.7 | 73.9±3.4 | 213.7±10.8 | 7.62±0.07 | 45.4 |
| Rochester (1995)[64] | GCGUUCGC<br>CGCUUGCG | 70.1±3.8 | 200.7±11.9 | 7.88±0.17 | 46.9 | 65.5±0.9 | 186.5±2.9 | 7.69±0.03 | 46.7 |
| % Uncertainty[b] | | | | | | 9.32% | 10.5% | 1.41% | |

[a] Tm calculated for $10^{-4}$ M oligomer concentration.

[b] % uncertainty is σ/mean, where σ is the standard deviation. This was calculated for each duplex. The mean of these three % uncertainties for $\Delta G^\circ_{37}$ is 2.1%.

Table S2. A-form helices used to fit nearest neighbor parameters for 1mΨ-A and 1mΨ-G base pair stacks. These experiments were performed at Poznan.

| Duplex | Average of curve fits | | | | $T_M^{-1}$ vs log $C_T$ plots | | | |
|---|---|---|---|---|---|---|---|---|
| | -ΔH° (kcal/mol) | -ΔS° (eu) | -ΔG°$_{37}$ (kcal/mol) | $T_M$† (°C) | -ΔH° (kcal/mol) | -ΔS° (eu) | -ΔG°$_{37}$ (kcal/mol) | $T_M$† (°C) |
| 1AGCGC<br>AUCGCG | 51.8±6.0 | 139.2±18.8 | 8.57±0.17 | 50.1 | 48.7±2.7 | 129.8±8.4 | 8.46±0.07 | 49.7 |
| 1CGCGC<br>AGCGCG | 57.5±2.9 | 152.9±9.0 | 10.04±0.14 | 57.4 | 59.3±1.3 | 158.4±3.9 | 10.13±0.07 | 57.3 |
| 1GGCCG<br>GCCGG1 | 50.6±2.0 | 134.7±6.1 | 8.80±0.15 | 57.6 | 50.6±3.1 | 134.8±9.5 | 8.79±0.19 | 57.4 |
| 1GGCGC<br>ACCGCG | 63.2±4.2 | 168.8±12.9 | 10.89±0.26 | 59.7 | 62.0±4.5 | 165.0±13.7 | 10.80±0.26 | 60.1 |
| 1UGCAG<br>GACGU1 | 35.2±2.4 | 97.6±8.0 | 4.90±0.10 | 30.6 | 37.9±1.7 | 106.9±5.5 | 4.80±0.06 | 29.6 |
| 1UGCGC<br>AACGCG | 51.2±4.9 | 133.7±14.5 | 9.72±0.41 | 57.7 | 46.7±4.7 | 120.2±14.3 | 9.43±0.31 | 57.5 |
| AAGCGC‡<br>1UCGCG | 45.9±6.1 | 120.9±18.7 | 8.41±0.33 | 50.2 | 38.6±1.1 | 98.2±3.5 | 8.14±0.04 | 50.5 |
| ACGCGC<br>1GCGCG | 59.6±5.7 | 158.9±17.4 | 10.30±0.34 | 58.0 | 56.9±6.6 | 150.9±20.0 | 10.13±0.38 | 57.7 |
| AGGCGC<br>1CCGCG | 58.3±7.1 | 154.7±21.3 | 10.36±0.46 | 58.6 | 54.4±2.4 | 143.1±7.3 | 10.06±0.14 | 58.2 |
| AUGCGC<br>1ACGCG | 57.1±6.5 | 151.0±19.6 | 10.427±0.48 | 58.7 | 50.8±3.9 | 131.9±11.8 | 9.91±0.23 | 59.0 |
| CC1AGG<br>GGA1CC | 56.4±2.7 | 154.5±8.4 | 8.45±0.12 | 53.2 | 57.4±0.8 | 157.7±2.6 | 8.48±0.02 | 53.0 |
| CCG1GG<br>GG1GCC | 50.8±2.8 | 147.1±8.9 | 5.19±0.06 | 34.0 | 56.9±0.7 | 166.9±6.2 | 5.09±0.04 | 34.1 |
| CG1AGC<br>GCA1CG | 60.6±5.6 | 167.1±17.3 | 8.72±0.33 | 48.9 | 59.4±6.0 | 163.6±19.2 | 8.70±0.22 | 48.5 |
| CGA1GC<br>GC1ACG | 50.5±3.8 | 139.7±12.1 | 7.20±0.13 | 41.0 | 45.8±1.1 | 121.9±6.4 | 7.11±0.02 | 47.2 |
| GCA1GC<br>CG1ACG | 66.7±5.6 | 185.2±17.0 | 9.26±0.31 | 54.6 | 65.4±4.9 | 181.2±15.1 | 9.16±0.26 | 54.7 |
| GCG1GC<br>CG1GCG | 46.1±1.1 | 129.3±3.6 | 6.31±0.12 | 39.2 | 40.9±2.6 | 111.5±8.2 | 6.29±0.06 | 41.9 |
| GG1ACC<br>CCA1GG | 59.3±3.9 | 161.1±12.1 | 9.37±0.20 | 57.4 | 59.8±1.9 | 162.6±5.0 | 9.40±0.11 | 57.4 |
| GGCGC1<br>1CGCGG | 57.0±1.6 | 155.5±4.8 | 8.80±0.19 | 54.8 | 54.6±2.8 | 147.9±8.8 | 8.68±0.14 | 55.4 |
| UC1AGA<br>AGA1CU | 45.8±2.9 | 130.6±9.4 | 5.29±0.10 | 34.4 | 47.8±2.1 | 137.3±6.8 | 5.23±0.05 | 34.0 |
| C1GC1GG‡<br>GAUGAUC | 53.1±5.6 | 148.6±17.7 | 7.07±0.26 | 39.8 | 45.5±8.3 | 124.1±26.9 | 7.05±0.47 | 40.3 |
| C1GC1GG‡<br>GG1CG1C | 41.2±2.1 | 113.9±6.7 | 5.88±0.08 | 38.5 | 45.1±2.9 | 126.9±9.5 | 5.86±0.06 | 37.5 |

| | | | | | | | | |
|---|---|---|---|---|---|---|---|---|
| CG11ACG‡<br>GCAAUGC | 60.9±3.6 | 167.5±10.5 | 8.96±0.31 | 49.8 | 49.1±1.0 | 130.6±3.1 | 8.57±0.03 | 50.6 |
| CGA1ACG<br>GC1A1GC | 59.4±1.4 | 160.2±4.4 | 9.69±0.09 | 54.6 | 62.3±2.3 | 169.1±7.1 | 9.81±0.10 | 54.5 |
| CGA1ACG<br>GCUAUGC | 59.6±3.3 | 164.8±10.0 | 8.51±0.18 | 47.5 | 52.0±1.4 | 141.1±4.5 | 8.27±0.04 | 47.5 |
| CGA1CCG<br>GCUAGGC | 64.7±2.1 | 174.7±6.5 | 10.52±0.10 | 57.4 | 67.7±4.5 | 183.8±13.8 | 10.66±0.25 | 57.3 |
| CGA1GCG<br>GCUACGC | 65.1±4.4 | 175.8±13.7 | 10.58±0.23 | 57.5 | 58.5±3.3 | 155.7±10.2 | 10.23±0.19 | 57.8 |
| CGA1UCG<br>GCUAAGC | 55.4±2.2 | 152.3±6.7 | 8.20±0.14 | 46.4 | 49.4±1.7 | 133.2±5.4 | 8.09±0.03 | 47.1 |
| CGC1ACG<br>GCGAUGC | 64.3±3.1 | 173.7±9.2 | 10.38±0.22 | 57.0 | 60.3±6.0 | 161.7±18.5 | 10.18±0.33 | 56.8 |
| CGC1CCG<br>GCGAGGC | 68.0±2.1 | 179.6±6.0 | 12.34±0.27 | 65.7 | 62.3±4.1 | 162.7±12.1 | 11.87±0.34 | 65.9 |
| CGC1GCG<br>GCGACGC | 70.6±3.1 | 186.9±9.2 | 12.63±0.24 | 66.3 | 76.3±3.4 | 203.8±10.2 | 13.05±0.27 | 66.2 |
| CGC1UCG<br>GCGAAGC | 63.7±3.2 | 172.5±9.9 | 10.17±0.16 | 56.0 | 65.4±2.0 | 177.8±6.0 | 10.22±0.10 | 55.7 |
| CGG1UCG<br>GCCAAGC | 65.2±3.8 | 176.2±11.8 | 10.54±0.21 | 57.4 | 59.0±4.8 | 157.3±14.7 | 10.23±0.26 | 57.6 |
| CGG1GCG<br>GCCACGC | 71.1±6.4 | 188.4±18.8 | 12.66±0.61 | 66.3 | 77.9±5.7 | 208.7±17.0 | 13.22±0.46 | 65.9 |
| CGG1CCG<br>GCCAGGC | 68.7±1.4 | 180.7±4.6 | 12.68±0.14 | 67.4 | 65.1±8.6 | 169.8±25.3 | 12.38±0.75 | 67.9 |
| CGG1ACG<br>GCCAUGC | 70.5±2.2 | 191.6±6.5 | 11.05±0.19 | 58.4 | 61.6±2.4 | 164.5±7.2 | 10.55±0.13 | 58.8 |
| CGU1ACG<br>GCAAUGC | 59.5±2.9 | 165.6±8.9 | 8.13±0.19 | 45.6 | 52.0±1.7 | 141.9±5.3 | 7.97±0.03 | 46.0 |
| CGU1CCG<br>GCAAGGC | 61.3±3.1 | 166.3±9.4 | 9.77±0.16 | 54.0 | 60.2±1.9 | 162.9±5.8 | 9.70±0.08 | 54.1 |
| CGU1GCG<br>GCAACGC | 64.1±3.0 | 173.8±9.3 | 10.16±0.18 | 55.8 | 64.7±3.7 | 175.8±11.4 | 10.17±0.19 | 55.5 |
| CGU1UCG<br>GCAAAGC | 55.0±2.6 | 152.6±8.2 | 7.73±0.11 | 43.6 | 55.6±1.6 | 144.9±5.1 | 7.68±0.02 | 61.9 |
| CUCGCUC<br>GAG1GAG | 72.7±6.4 | 204.1±19.5 | 9.40±0.34 | 49.7 | 66.8±5.7 | 185.8±17.7 | 9.15±0.23 | 49.8 |
| AC11AAGU<br>UGAA11CA | 54.2±4.6 | 151.5±14.4 | 7.23±0.17 | 46.0 | 49.1±2.3 | 133.5±7.4 | 7.09±0.05 | 50.3 |
| AGA1A1CU<br>UC1A1AGA | 61.6±3.2 | 169.9±9.8 | 8.88±0.23 | 54.2 | 53.3±2.9 | 144.5±9.0 | 8.53±0.12 | 54.2 |
| C11CGUUC<br>GAAGUAAG | 63.5±0.7 | 180.0±2.4 | 6.79±0.06 | 42.7 | 68.0±2.6 | 197.4±8.4 | 6.77±0.03 | 38.1 |
| C1A1A1AG<br>GA1A1A1C | 65.1±4.1 | 181.4±12.4 | 8.88±0.28 | 52.8 | 57.8±2.7 | 158.7±8.2 | 8.57±0.10 | 53.4 |
| C1ACG1AG | 74.5±2.1 | 203.0±6.1 | 11.56±0.22 | 63.5 | 65.0±1.7 | 174.6±5.2 | 10.90±0.12 | 63.8 |

| | | | | | | | | |
|---|---|---|---|---|---|---|---|---|
| GA1GCA1C | | | | | | | | |
| C1A1GA1G<br>GA1AU1AC | 63.7±2.7 | 183.3±9.1 | 6.85±0.12 | 38.6 | 67.1±3.5 | 194.3±11.4 | 6.79±0.05 | 38.4 |
| C1GUA1AG<br>GA1AUG1C | 55.9±2.5 | 165.6±8.5 | 4.51±0.11 | 30.8 | 58.4±1.0 | 174.0±3.6 | 4.40±0.04 | 30.5 |
| C1G1A1AG<br>GA1A1G1C | 54.6±1.4 | 156.7±4.6 | 5.96±0.13 | 34.0 | 56.6±4.1 | 163.4±13.2 | 5.94±0.09 | 33.7 |
| C1GC1AGC‡<br>GAUGA11G | 49.2±7.4 | 141.2±24.8 | 5.42±0.26 | 30.1 | 55.7±3.3 | 162.9±10.9 | 5.17±0.11 | 29.6 |
| CCAG11GG<br>GG11GACC | 70.7±6.1 | 200.4±19.0 | 8.55±0.29 | 50.1 | 64.9±5.9 | 182.4±18.5 | 8.32±0.22 | 50.2 |
| CCUG1AGG<br>GGA1GUCC | 73.2±6.3 | 208.5±19.6 | 8.57±0.29 | 49.6 | 65.0±6.7 | 183.1±20.7 | 8.26±0.27 | 49.6 |
| CG1UGUAG<br>GCAG1G1C | 67.0±3.3 | 197.0±10.8 | 5.91±0.09 | 34.1 | 73.3±4.0 | 217.8±13.2 | 5.78±0.10 | 33.7 |
| CUCGUGAG‡<br>GAG1GCUC | 54.8±1.3 | 152.1±4.4 | 7.59±0.11 | 43.3 | 66.2±1.8 | 188.8±5.6 | 7.70±0.02 | 42.3 |
| CUAG1GAG<br>GA11G1UC | 63.5±2.1 | 187.4±7.1 | 5.36±0.12 | 31.5 | 73.2±1.0 | 219.8±2.6 | 5.08±0.04 | 30.8 |
| CUCG1GAG<br>GAG1GCUC | 68.1±2.9 | 194.0±9.0 | 7.96±0.13 | 47.6 | 64.7±3.0 | 183.4±9.4 | 7.85±0.08 | 47.6 |
| CUC1GCUC<br>GAGG1GAG | 90.7±8.7 | 256.0±26.8 | 11.34±0.37 | 54.2 | 84.5±3.2 | 237.0±9.9 | 11.00±0.14 | 54.3 |
| CUCG1CUC<br>GAG1GGAG | 77.6±5.7 | 219.2±17.7 | 9.64±0.23 | 49.8 | 72.5±3.1 | 203.2±9.6 | 9.47±0.10 | 50.1 |
| CUCG1CUC<br>GAGUGGAG | 69.9±3.7 | 198.9±11.7 | 8.25±0.15 | 44.6 | 72.0±2.8 | 205.3±9.0 | 8.29±0.06 | 44.9 |
| CUCGGCUC<br>GAG11GAG | 73.5±3.6 | 205.9±11.2 | 9.63±0.14 | 50.7 | 73.3±5.6 | 205.4±17.5 | 9.59±0.18 | 50.5 |
| CUCGGCUC<br>GAG1UGAG | 71.7±3.2 | 203.0±10.3 | 8.70±0.10 | 46.9 | 71.0±3.1 | 201.0±9.9 | 8.66±0.07 | 46.6 |
| CUCGGCUC<br>GAGU1GAG | 75.6±3.0 | 213.8±9.3 | 9.28±0.20 | 48.8 | 67.1±5.2 | 187.2±16.4 | 9.06±0.17 | 49.1 |
| CUCUGCUC<br>GAGG1GAG | 94.6±3.1 | 269.6±9.6 | 10.97±0.14 | 52.3 | 101.3±7.8 | 290.6±24.1 | 11.22±0.31 | 51.9 |
| CUU1G1UC<br>GAAGUAAG | 63.4±2.0 | 186.1±5.1 | 5.66±0.04 | 32.9 | 70.7±3.7 | 210.2±12.2 | 5.47±0.10 | 32.6 |
| CUU1G1UC<br>GAAG1AAG | 70.1±3.0 | 206.3±9.7 | 6.08±0.14 | 35.2 | 67.0±4.2 | 196.2±13.7 | 6.14±0.09 | 35.2 |
| CUUG1UUC<br>GAAUGAAG | 68.4±3.8 | 208.8±12.9 | 3.61±0.20 | 24.4 | 66.7±1.8 | 203.3±6.2 | 3.67±0.10 | 24.1 |
| CUUGAU1C<br>GAAU1GAG | 65.2±3.1 | 196.7±10.6 | 4.21±0.19 | 26.3 | 72.4±0.9 | 221.1±3.1 | 3.88±0.04 | 25.8 |
| CUUGAU1C | 61.0±2.9 | 179.5±9.6 | 5.35±0.10 | 31.0 | 64.8±2.3 | 194.9±7.4 | 5.23±0.06 | 26.9 |

| | | | | | | | | |
|---|---|---|---|---|---|---|---|---|
| GAA11AAG | | | | | | | | |
| CUUGGUUC<br>GAA11AAG | 65.1±2.9 | 193.7±9.8 | 5.01±0.16 | 30.0 | 65.8±2.8 | 196.2±9.5 | 4.96±0.10 | 29.7 |
| G11A1GGC‡<br>CAA1A1UG | 65.5±6.2 | 182.7±19.2 | 8.88±0.40 | 48.3 | 48.5±4.9 | 129.0±15.3 | 8.45±0.24 | 50.1 |
| G11A1GGC<br>CGG1A11G | 55.5±9.5 | 165.6±32.0 | 4.13±0.40 | 28.6 | 53.9±5.5 | 160.5±18.6 | 4.11±0.28 | 28.3 |
| GACGCG11<br>11GCGCAG | 68.3±3.4 | 186.1±10.3 | 10.59±0.23 | 61.0 | 60.9±6.1 | 163.6±18.3 | 10.13±0.43 | 61.6 |
| GCAGCUG1<br>1GUCGACG | 76.7±1.3 | 210.7±3.7 | 11.32±0.20 | 61.8 | 83.3±8.7 | 230.6±26.0 | 11.75±0.60 | 61.5 |
| GCAG1UGC<br>CGU1GACG | 71.2±3.0 | 202.4±9.1 | 8.44±0.20 | 49.5 | 67.6±5.4 | 191.2±16.9 | 8.33±0.17 | 49.5 |
| GCUGG1GC<br>CGAU1ACG | 72.9±2.7 | 204.2±8.3 | 9.55±0.12 | 50.5 | 77.0±5.1 | 217.2±16.1 | 9.66±0.17 | 50.0 |
| GCUGG1GC<br>CGAUUACG | 63.4±3.7 | 176.6±12.0 | 8.65±0.06 | 47.6 | 63.7±7.5 | 177.3±23.6 | 8.65±0.28 | 48.0 |
| GCUGGUGC<br>CGAU1ACG | 77.4±11.0 | 220.4±34.5 | 9.10±0.36 | 47.4 | 72.4±4.8 | 204.9±15.3 | 8.90±0.12 | 47.3 |
| GCUGGUGC‡<br>CGA11ACG | 78.4±7.3 | 223.2±22.2 | 9.16±0.43 | 47.8 | 61.0±1.4 | 168.9±4.4 | 8.64±0.04 | 48.0 |
| GCUGGUGC<br>CGA1UACG | 71.3±3.0 | 203.4±9.9 | 8.19±0.11 | 44.5 | 73.6±3.8 | 210.9±11.9 | 8.21±0.08 | 44.2 |
| GG11A1GG<br>CCAG1G1U | 52.4±3.4 | 147.6±10.9 | 6.64±0.03 | 37.5 | 57.1±1.7 | 162.9±5.4 | 6.60±0.02 | 37.3 |
| GG1UGACC<br>CCAGU1GG | 84.1±3.2 | 236.6±9.8 | 10.74±0.17 | 56.8 | 87.0±5.5 | 245.5±16.7 | 10.89±0.31 | 56.6 |
| GGA1GUCC<br>CCUG1AGG | 80.8±5.0 | 228.3±15.3 | 9.99±0.27 | 54.5 | 80.6±6.0 | 227.7±18.5 | 9.94±0.30 | 54.5 |
| GGC11CAA<br>CCGAAGUU | 88.7±7.2 | 246.7±21.8 | 13.15±0.45 | 58.1 | 85.8±7.6 | 234.7±22.8 | 12.98±0.52 | 62.3 |
| GU1A1AAC<br>CAA1A1UG | 57.7±3.5 | 162.2±11.4 | 7.38±0.21 | 46.5 | 65.6±8.8 | 187.5±28.1 | 7.41±0.26 | 45.6 |
| U1ACGUAG<br>GAUGCA1U | 56.2±2.3 | 158.0±7.3 | 7.25±0.13 | 45.6 | 52.2±2.7 | 145.1±8.7 | 7.15±0.07 | 46.3 |
| CG11A1ACG‡<br>GCAA1A1GC | 80.9±0.4 | 218.4±2.8 | 13.72±0.28 | 64.7 | 66.5±9.2 | 175.1±27.3 | 12.15±0.75 | 65.9 |
| CG11GAAGC<br>GCGAC11CG | 84.8±1.9 | 231.7±5.7 | 13.93±0.18 | 62.4 | 85.7±4.9 | 234.4±14.9 | 12.99±0.34 | 62.3 |
| CGAAG11GC‡<br>GC11CAACG | 84.4±2.0 | 227.1±6.4 | 14.01±0.23 | 67.0 | 68.4±8.3 | 179.4±17.8 | 12.76±0.49 | 68.1 |
| AGGA1G11GG<br>UCCUGCGACC | 95.1±1.0 | 260.3±3.3 | 14.34±0.04 | 64.9 | 88.3±6.3 | 240.2±19.0 | 13.85±0.47 | 64.8 |
| AGGA1GU1GG<br>UCCUGCGACC | 102.8±7.7 | 287.2±23.6 | 13.74±0.46 | 60.3 | 101.5±10.4 | 283.2±31.3 | 13.66±0.70 | 60.5 |
| | 70.5±4.6 | 210.9±14.9 | 5.05±0.06 | 30.8 | 84.9±3.5 | 258.9±11.8 | 4.62±0.12 | 30.1 |

| | | | | | | | | |
|---|---|---|---|---|---|---|---|---|
| C1GGUG11AU‡<br>GG11AUAG1G | | | | | | | | |
| CAGAGGAGAC<br>GUCUU1UCUG | 115.1±11.9 | 333.5±37.6 | 11.66±0.10 | 51.5 | 106.7±8.9 | 307.3±27.7 | 11.36±0.27 | 51.8 |
| CUCUU1ACUC<br>GAGAGG1GAG | 97.8±7.6 | 278.6±23.4 | 11.40±0.37 | 53.2 | 95.8±5.6 | 272.5±17.4 | 11.26±0.24 | 53.2 |
| CUCUUUACUC‡<br>GAGAGG1GAG | 65.4±1.6 | 190.8±5.4 | 6.25±0.09 | 35.6 | 77.8±6.6 | 231.8±22.0 | 5.88±0.20 | 34.5 |
| G1GAAUUUAC<br>CAUUUAAG1G | Very low temperature melting | | | | | | | |

†$T_M$ calculated for $10^{-4}$ M oligomer concentration.

‡The ΔH° values as fit by the two methods differed by greater than 15%. This can indicate non-two-state melting. These duplexes were excluded from the fit.

Table S3. The helical stack nearest neighbor parameters for stacks with 1mΨ-A base pairs. The character "1" is used to represent 1mΨ. σ, the uncertainty, is the standard error of the regression. The U analog values are from Xia et al. [13]. The stack parameters are read as one pair following another from left to right. Therefore, the top row in a stack is a strand from 5' to 3', and the bottom row is a strand from 3' to 5'.

| 1mΨ-A stack: | $\Delta G°_{37}$ (kcal/mol): | σ: | analog: | $\Delta G°_{37}$ (kcal/mol): | σ: | # Occurrences in dataset |
|---|---|---|---|---|---|---|
| AC<br>1G | -2.80 | 0.24 | AC<br>UG | -2.24 | 0.06 | 18 |
| 1G<br>AC | -2.58 | 0.25 | UG<br>AC | -2.11 | 0.07 | 16 |
| 1C<br>AG | -2.53 | 0.25 | UC<br>AG | -2.35 | 0.06 | 14 |
| AG<br>1C | -2.22 | 0.24 | AG<br>UC | -2.08 | 0.06 | 25 |
| 1A<br>A1 | -1.81 | 0.45 | UA<br>AU | -1.33 | 0.09 | 16 |
| 1A<br>AU | -1.67 | 0.27 | UA<br>AU | -1.33 | 0.09 | 8 |
| A1<br>UA | -1.65 | 0.28 | AU<br>UA | -1.10 | 0.08 | 6 |
| A1<br>1A | -1.51 | 0.46 | AU<br>UA | -1.10 | 0.08 | 11 |
| 1U<br>AA | -1.41 | 0.28 | UU<br>AA | -0.93 | 0.03 | 7 |
| 11<br>AA | -1.23 | 0.25 | UU<br>AA | -0.93 | 0.03 | 5 |
| AA<br>1U | -1.12 | 0.28 | AA<br>UU | -0.93 | 0.03 | 7 |

Table S4: The helical stack nearest neighbor parameters for stacks with 1mΨ-G base pairs or with 1mΨ-A and an adjacent U-G pair. The character "1" is used to represent 1mΨ. σ, the uncertainty, is the standard error of the regression. The U analog values are from Chen et al.[30].

| 1mΨ-G stack: | $\Delta G^\circ_{37}$ (kcal/mol): | σ: | analog: | $\Delta G^\circ_{37}$ (kcal/mol): | σ: | # Occurrences in dataset |
|---|---|---|---|---|---|---|
| 1C<br>GG | -2.63 | 0.47 | GG<br>CU | -1.80 | 0.09 | 2 |
| G1<br>CG | -2.25 | 0.21 | GU<br>CG | -2.15 | 0.10 | 11 |
| C1<br>GG | -2.01 | 0.25 | CU<br>GG | -1.77 | 0.09 | 3 |
| 1G<br>GU | -2.00 | 0.43 | UG<br>GU | -0.57 | 0.19 | 3 |
| 1G<br>G1 | -1.94 | 0.42 | UG<br>GU | -0.57 | 0.19 | 5 |
| 1G<br>GC | -1.67 | 0.18 | CG<br>GU | -1.25 | 0.09 | 14 |
| 1U<br>AG | -1.32 | 0.34 | UU<br>AG | -0.51 | 0.08 | 5 |
| A1<br>1G | -1.16 | 0.39 | AU<br>UG | -0.90 | 0.08 | 5 |
| 1U<br>GG | -0.91 | 0.34 | GG<br>UU | -0.25 | 0.16 | 4 |
| 1G<br>A1 | -0.88 | 0.35 | UG<br>AU | -0.39 | 0.09 | 8 |
| 1A<br>GU | -0.87 | 0.28 | UG<br>AU | -0.39 | 0.09 | 5 |
| 1U<br>GA | -0.86 | 0.21 | AG<br>UU | -0.35 | 0.08 | 5 |
| A1<br>UG | -0.67 | 0.24 | AU<br>UG | -0.90 | 0.08 | 8 |
| 11<br>GG | -0.64 | 0.31 | GG<br>UU | -0.25 | 0.16 | 4 |
| GA<br>1U | -0.64 | 0.41 | UU<br>AG | -0.51 | 0.08 | 4 |
| AG<br>1U | -0.59 | 0.33 | AG<br>UU | -0.35 | 0.08 | 4 |
| 11<br>GA | -0.44 | 0.37 | AG<br>UU | -0.35 | 0.08 | 5 |
| 11<br>AG | -0.44 | 0.33 | UU<br>AG | -0.51 | 0.08 | 4 |
| AU<br>1G | -0.29 | 0.43 | AU<br>UG | -0.90 | 0.08 | 3 |
| G1<br>1G | -0.25 | 0.38 | GU<br>UG | 0.72 | 0.19 | 8 |

| | | | | | | |
|---|---|---|---|---|---|---|
| GG<br>1U | 0.00 | 0.35 | GG<br>UU | -0.25 | 0.16 | 4 |
| GU<br>1G | 0.21 | 0.43 | GU<br>UG | 0.72 | 0.19 | 3 |
| 1G<br>AU | 0.34 | 0.63 | UG<br>AU | -0.39 | 0.09 | 2 |

Table S5. The agreement between measured and estimated folding stability for duplexes composed of canonical base pairs. Location is either IBCh (Institute of Bioorganic Chemistry) or DS (DNA Software). The uncertainties for the experiment values, σ, are 4% of the experimental value[13]. The uncertainties for the estimated values are propagated from nearest neighbor parameters. The character "1" is used to represent 1mΨ. The residual is the difference in $\Delta G°_{37}$ between the experiment and the estimate.

| Sequence | Location | Experimental $-\Delta G°_{37}$ (kcal/mol) | Σ (kcal/mol) | Estimated $-\Delta G°_{37}$ (kcal/mol) | Σ (kcal/mol) | Residual (kcal/mol) |
|---|---|---|---|---|---|---|
| 1GGCCG<br>GCCGG1 | IBCh | 8.79 | 0.35 | 8.76 | 0.45 | -0.03 |
| 1UGCAG<br>GACGU1 | IBCh | 4.80 | 0.19 | 4.84 | 0.50 | 0.04 |
| 1AGCGC<br>AUCGCG | IBCh | 8.46 | 0.34 | 8.86 | 0.40 | 0.40 |
| 1CGCGC<br>AGCGCG | IBCh | 10.13 | 0.41 | 10.00 | 0.41 | -0.13 |
| 1GGCGC<br>ACCGCG | IBCh | 10.80 | 0.43 | 10.95 | 0.39 | 0.15 |
| 1UGCGC<br>AACGCG | IBCh | 9.43 | 0.38 | 8.63 | 0.41 | -0.80 |
| 1CGCGA<br>AGCGC1 | DS | 7.15 | 0.29 | 8.68 | 0.58 | 1.53 |
| ACGCG1<br>1GCGCA‡ | DS | 6.80 | 0.27 | 9.22 | 0.56 | 2.42 |
| CCG1GG<br>GG1GCC | IBCh | 5.09 | 0.20 | 5.59 | 0.58 | 0.50 |
| CC1AGG<br>GGA1CC | IBCh | 8.48 | 0.34 | 8.25 | 0.71 | -0.23 |
| GGCGC1<br>1CGCGG | IBCh | 8.68 | 0.35 | 8.70 | 0.58 | 0.02 |
| GCG1GC<br>CG1GCG | IBCh | 6.29 | 0.25 | 5.91 | 0.59 | -0.38 |
| GCA1GC<br>CG1ACG | IBCh | 9.16 | 0.37 | 8.99 | 0.73 | -0.17 |
| GG1ACC<br>CCA1GG | IBCh | 9.40 | 0.38 | 9.41 | 0.71 | 0.01 |

| | | | | | | |
|---|---|---|---|---|---|---|
| GCGCC1<br>CGCGGA | IBCh | 10.06 | 0.40 | 10.59 | 0.38 | 0.53 |
| GCGCG1<br>CGCGCA | IBCh | 10.13 | 0.41 | 10.27 | 0.40 | 0.14 |
| GC1ACG<br>CGA1GC | IBCh | 8.70 | 0.35 | 8.52 | 0.62 | -0.18 |
| GCA1CG<br>CG1AGC | IBCh | 7.11 | 0.28 | 8.31 | 0.63 | 1.20 |
| GCGCA1<br>CGCGUA | IBCh | 9.91 | 0.40 | 8.87 | 0.41 | -1.04 |
| UC1AGA<br>AGA1CU | IBCh | 5.23 | 0.21 | 5.53 | 0.71 | 0.30 |
| CUCGCUC<br>GAG1GAG | IBCh | 9.15 | 0.37 | 8.69 | 0.39 | -0.46 |
| CGA1ACG<br>GC1A1GC | IBCh | 9.81 | 0.39 | 9.28 | 0.78 | -0.53 |
| CGA1ACG<br>GCUAUGC | IBCh | 8.27 | 0.33 | 8.54 | 0.49 | 0.27 |
| CGC1ACG<br>GCGAUGC | IBCh | 10.18 | 0.41 | 10.18 | 0.47 | 0.00 |
| CGG1ACG<br>GCCAUGC | IBCh | 10.55 | 0.42 | 10.60 | 0.47 | 0.05 |
| CGU1ACG<br>GCAAUGC | IBCh | 7.97 | 0.32 | 7.90 | 0.50 | -0.07 |
| CGA1CCG<br>GCUAGGC | IBCh | 10.66 | 0.43 | 10.42 | 0.48 | -0.24 |
| CGC1CCG<br>GCGAGGC | IBCh | 11.87 | 0.47 | 12.06 | 0.46 | 0.19 |
| CGG1CCG<br>GCCAGGC | IBCh | 12.38 | 0.50 | 12.48 | 0.47 | 0.10 |
| CGU1CCG<br>GCAAGGC | IBCh | 9.70 | 0.39 | 9.78 | 0.48 | 0.08 |
| CGA1GCG<br>GCUACGC | IBCh | 10.23 | 0.41 | 10.63 | 0.48 | 0.40 |
| CGC1GCG<br>GCGACGC | IBCh | 13.05 | 0.52 | 12.27 | 0.48 | -0.78 |
| CGG1GCG | IBCh | 13.22 | 0.53 | 12.69 | 0.46 | -0.53 |

| | | | | | | |
|---|---|---|---|---|---|---|
| GCCACGC | | | | | | |
| CGU1GCG GCAACGC | IBCh | 10.17 | 0.41 | 9.99 | 0.48 | -0.18 |
| CGA1UCG GCUAAGC | IBCh | 8.09 | 0.32 | 8.39 | 0.50 | 0.30 |
| CGC1UCG GCGAAGC | IBCh | 10.22 | 0.41 | 10.03 | 0.48 | -0.19 |
| CGG1UCG GCCAAGC | IBCh | 10.23 | 0.41 | 10.45 | 0.47 | 0.22 |
| CGU1UCG GCAAAGC | IBCh | 7.68 | 0.31 | 7.75 | 0.49 | 0.07 |
| 11CGCGAG GAGCGC11† | DS | 9.42 | 0.38 | 9.56 | 0.94 | 0.14 |
| 1ACGCG1G G1GCGCA1 | DS | 10.60 | 0.42 | 10.98 | 0.90 | 0.38 |
| 1GACGUCG GCUGCAG1 | DS | 9.91 | 0.40 | 10.36 | 0.46 | 0.45 |
| 1CUGCAGG GGACGUC1‡ | DS | 9.44 | 0.38 | 12.54 | 0.99 | 3.10 |
| AC11AAGU UGAA11CA | IBCh | 7.09 | 0.28 | 7.77 | 0.87 | 0.68 |
| AGA1A1CU UC1A1AGA | IBCh | 8.53 | 0.34 | 8.63 | 1.17 | 0.10 |
| CCUG1AGG GGA1GUCC | IBCh | 8.26 | 0.33 | 8.15 | 0.74 | -0.11 |
| CUCGGCUC GAG11GAG | IBCh | 9.59 | 0.38 | 9.33 | 0.50 | -0.26 |
| CUCGGCUC GAG1UGAG | IBCh | 8.66 | 0.35 | 8.59 | 0.49 | -0.07 |
| CUCGGCUC GAGU1GAG | IBCh | 9.06 | 0.36 | 9.18 | 0.49 | 0.12 |
| CUCG1GAG GAG1GCUC | IBCh | 7.85 | 0.31 | 7.93 | 0.59 | 0.08 |
| CCAG11GG GG11GACC | IBCh | 8.32 | 0.33 | 8.29 | 1.00 | -0.03 |
| C1G1A1AG GA1A1G1C | IBCh | 5.94 | 0.24 | 5.81 | 1.26 | -0.13 |

| | | | | | | |
|---|---|---|---|---|---|---|
| C1GUA1AG<br>GA1AUG1C | IBCh | 4.40 | 0.18 | 4.35 | 1.00 | -0.05 |
| CG1UGUAG<br>GCAG1G1C | IBCh | 5.78 | 0.23 | 6.06 | 1.03 | 0.28 |
| CUAG1GAG<br>GA11G1UC | IBCh | 5.08 | 0.20 | 5.01 | 0.87 | -0.07 |
| C1A1GA1G<br>GA1AU1AC | IBCh | 6.79 | 0.27 | 6.52 | 1.32 | -0.27 |
| C1ACG1AG<br>GA1GCA1C | IBCh | 10.9 | 0.44 | 11.50 | 1.15 | 0.60 |
| C1A1A1AG<br>GA1A1A1C | IBCh | 8.57 | 0.34 | 8.37 | 1.72 | -0.20 |
| GGC11CAA<br>CCGAAGUU | IBCh | 12.98 | 0.52 | 11.16 | 0.50 | -1.82 |
| GAGG1GAG<br>CUC1GCUC | IBCh | 9.47 | 0.38 | 9.32 | 0.69 | -0.15 |
| GAGGUGAG<br>CUC1GCUC | IBCh | 8.29 | 0.33 | 8.44 | 0.70 | 0.15 |
| GACGCG11<br>11GCGCAG | IBCh | 10.13 | 0.41 | 10.10 | 0.87 | -0.03 |
| GG1UGACC<br>CCAGU1GG | IBCh | 10.89 | 0.44 | 10.81 | 0.89 | -0.08 |
| GAG1GGAG<br>CUCG1CUC | IBCh | 11.00 | 0.44 | 10.97 | 0.60 | -0.03 |
| GAG1GGAG<br>CUCGUCUC | IBCh | 11.22 | 0.45 | 10.79 | 0.56 | -0.43 |
| GCUGG1GC<br>CGAU1ACG | IBCh | 9.66 | 0.39 | 9.87 | 0.64 | 0.21 |
| GCUGG1GC<br>CGAUUACG | IBCh | 8.65 | 0.35 | 8.34 | 0.60 | -0.31 |
| GCUGGUGC<br>CGAU1ACG | IBCh | 8.90 | 0.36 | 8.91 | 0.51 | 0.01 |
| GCAG1UGC<br>CGU1GACG | IBCh | 8.33 | 0.33 | 8.51 | 0.64 | 0.18 |
| GCAGCUG1<br>1GUCGACG | IBCh | 11.75 | 0.47 | 11.78 | 0.51 | 0.03 |
| GCUGGUGC | IBCh | 8.21 | 0.33 | 8.71 | 0.54 | 0.50 |

| | | | | | | |
|---|---|---|---|---|---|---|
| CGA1UACG | | | | | | |
| GGA1GUCC<br>CCUG1AGG | IBCh | 9.94 | 0.40 | 9.98 | 0.70 | 0.04 |
| GAA1GAAG<br>CU1G1UUC | IBCh | 6.14 | 0.25 | 6.42 | 0.79 | 0.28 |
| GAAUGAAG<br>CU1G1UUC | IBCh | 5.47 | 0.22 | 5.61 | 0.82 | 0.14 |
| GAA11AAG<br>CUUGGUUC | IBCh | 4.96 | 0.20 | 4.38 | 0.54 | -0.58 |
| GAAGUAAG<br>CUU1GUUC | IBCh | 3.67 | 0.15 | 3.24 | 0.54 | -0.43 |
| GAA11AAG<br>C1UAGUUC | IBCh | 5.23 | 0.21 | 5.53 | 0.68 | 0.30 |
| GAG1UAAG<br>C1UAGUUC | IBCh | 3.88 | 0.16 | 4.04 | 0.73 | 0.16 |
| GG11A1GG<br>CCAG1G1U | IBCh | 6.60 | 0.26 | 6.39 | 0.89 | -0.21 |
| G11A1GGC<br>CGG1A11G | IBCh | 4.11 | 0.16 | 4.53 | 1.14 | 0.42 |
| GU1A1AAC<br>CAA1A1UG | IBCh | 7.41 | 0.30 | 7.33 | 1.18 | -0.08 |
| GAAUGAAG<br>CUUGC11C | IBCh | 6.77 | 0.27 | 7.32 | 0.50 | 0.55 |
| GAC1AGUC<br>CUGA1CAG | DS | 10.64 | 0.43 | 10.91 | 0.71 | 0.27 |
| GACA1GUC<br>CUG1ACAG | DS | 10.90 | 0.44 | 11.33 | 0.73 | 0.43 |
| G1CGCGA1<br>1AGCGC1G | DS | 10.90 | 0.44 | 11.00 | 0.97 | 0.10 |
| GCUGCAG1<br>1GACGUCG | DS | 11.65 | 0.47 | 11.78 | 0.51 | 0.13 |
| GGACGUC1<br>1CUGCAGG | DS | 9.88 | 0.40 | 11.04 | 0.58 | 1.16 |
| GACGCG11<br>11GCGCAG† | DS | 9.64 | 0.39 | 10.10 | 0.87 | 0.46 |
| U1ACGUAG<br>GAUGCA1U | IBCh | 7.15 | 0.29 | 6.84 | 0.89 | -0.31 |

| | | | | | | |
|---|---|---|---|---|---|---|
| GC11CAGCG<br>CGAAG11GC | IBCh | 12.99 | 0.52 | 12.94 | 0.70 | -0.05 |
| CCAGCGUCCU<br>GG11G1AGGA | IBCh | 13.85 | 0.55 | 14.02 | 0.64 | 0.17 |
| CCAGCGUCCU<br>GG1UG1AGGA | IBCh | 13.66 | 0.55 | 14.07 | 0.59 | 0.41 |
| CAGAGGAGAC<br>GUCUU1UCUG | IBCh | 11.36 | 0.45 | 11.02 | 0.61 | -0.34 |
| GAG1GGAGAG<br>CUCA1UUCUC | IBCh | 11.26 | 0.45 | 11.04 | 0.64 | -0.22 |

†These duplexes are considered non-two-state because the deviation of ΔH° between the two fit methods is greater than 15%. They are included in the comparison.
‡These duplexes are outliers in terms of agreement between the experimental $\Delta G°_{37}$ and the estimate.

Table S6. Helical stabilities measured by DNA Software. The character “1” is used to represent 1mΨ.

| | Average of curve fits | | | | $T_M^{-1}$ vs log $C_T$ plots | | | |
|---|---|---|---|---|---|---|---|---|
| Duplex | -ΔH° (kcal/mol) | -ΔS° (eu) | -ΔG°$_{37}$ (kcal/mol) | $T_M$† (˚C) | -ΔH° (kcal/mol) | -ΔS° (eu) | -ΔG°$_{37}$ (kcal/mol) | $T_M$† (˚C) |
| 1CGCGA<br>AGCGC1 | 51.01±3.25 | 141.38±10.1 | 7.16±0.13 | 46.3 | 43.94±5.07 | 118.64±16.08 | 7.15±0.16 | 47.7 |
| ACGCG1<br>1GCGCA | 45.75±2.91 | 125.49±9.45 | 6.83±0.23 | 45 | 50.06±24.16 | 139.47±77.03 | 6.8±2.03 | 44.1 |
| 11CGCGAG<br>GAGCGC11 | 61.44±8.01 | 166.06±24.38 | 9.94±0.45 | 60.10 | 51.35±3.4 | 135.19±10.44 | 9.42±0.17 | 61.4 |
| 1ACGCG1G<br>G1GCGCA1 | 68.52±7.73 | 186.49±23.77 | 10.68±0.38 | 61.40 | 67.55±9.5 | 183.61±29 | 10.6±0.58 | 61.4 |
| 1CUGCAGG<br>GGACGUC1 | 67.51±5.5 | 186.35±17.18 | 9.71±0.24 | 56.70 | 60.87±7.79 | 165.81±24.08 | 9.44±0.39 | 57.5 |
| 1GACGUCG<br>GCUGCAG1 | 70.39±6.58 | 193.88±20.01 | 10.26±0.38 | 58.60 | 62.97±4.54 | 171.08±13.98 | 9.91±0.22 | 59.4 |
| G1CGCGA1<br>1AGCGC1G | 76.92±6.13 | 210.98±18.45 | 11.48±0.42 | 62.30 | 67.23±6.44 | 181.62±19.55 | 10.9±0.4 | 63.1 |
| GAC1AGUC<br>CUGA1CAG | 76.27±5.5 | 211.08±16.71 | 10.8±0.32 | 59.4 | 73.03±6.68 | 201.15±20.51 | 10.64±0.34 | 59.6 |
| GACA1GUC<br>CUG1ACAG | 78.33±5.23 | 216.93±15.96 | 11.05±0.28 | 59.8 | 75.8±3.02 | 209.25±9.25 | 10.9±0.15 | 60 |
| GACGCG11<br>11GCGCAG | 66.41±3.7 | 181.14±11.43 | 10.23±0.22 | 59.80 | 53.84±7.26 | 141.51±22.22 | 9.64±0.43 | 61.7 |
| GCUGCAG1<br>1GACGUCG | 85.08±7.91 | 236.65±24.08 | 11.68±0.45 | 60.60 | 85.08±3.77 | 236.76±11.49 | 11.65±0.21 | 60.4 |
| GGACGUC1<br>1CUGCAGG | 68.65±3.57 | 188.62±10.68 | 10.15±0.27 | 58.60 | 61.72±10.52 | 167.17±32.5 | 9.88±0.58 | 59.6 |

†$T_M$ is calculated for strand concentration of 0.1 mM.

Table S7. Dangling end model systems studied by optical melting. The character "1" is used to represent 1mΨ. Site is either the Institute of Bioorganic Chemistry (IBCh) or DNA Software (DS).

| | | **Average of curve fits** | | | | **$T_M^{-1}$ vs log $C_T$ plots** | | | |
|---|---|---|---|---|---|---|---|---|---|
| Site | Duplex | **-ΔH° (kcal/mol)** | **-ΔS° (eu)** | **-ΔG°$_{37}$ (kcal/mol)** | **$T_M$‡ (°C)** | **-ΔH° (kcal/mol)** | **-ΔS° (eu)** | **-ΔG°$_{37}$ (kcal/mol)** | **$T_M$‡ (°C)** |
| IBCh | 1AUGCAU<br>UACGUA1 | 42.5±2.1 | 119.3±6.8 | 5.51±0.07 | 35.7 | 46.7±1.6 | 133.0±5.2 | 5.47±0.03 | 35.5 |
| IBCh | AUGCAU1†<br>1UACGUA | 43.1±0.8 | 121.5±2.5 | 5.44±0.10 | 35.1 | 55.2±2.4 | 162.2±7.9 | 5.20±0.02 | 32.7 |
| IBCh | AUGCA11<br>11ACGUA | 41.9±1.8 | 116.9±6.1 | 5.67±0.14 | 36.8 | 46.6±2.3 | 133.4±7.5 | 5.60±0.05 | 34.0 |
| IBCh | AUGCA1G<br>G1ACGUA | 52.7±1.0 | 147.3±3.3 | 7.01±0.08 | 45.1 | 54.6±1.9 | 153.2±6.1 | 7.03±0.04 | 45.2 |
| IBCh | C1CAUGA<br>AGUAC1C | 50.3±1.9 | 144.0±6.2 | 5.66±0.11 | 36.8 | 55.6±2.5 | 161.5±6.8 | 5.58±0.05 | 36.1 |
| IBCh | 1GCGCAA<br>AACGCG1 | 59.7±3.3 | 159.3±9.7 | 10.27±0.25 | 63.0 | 60.6±3.5 | 162.1±10.5 | 10.35±0.24 | 62.8 |
| DS | ACGCG1A<br>A1GCGCA | 63.8±2.1 | 173.8±6.5 | 9.9±0.10 | 59.0 | 65.0±8.9 | 177.5±27.2 | 9.92±0.5 | 58.8 |
| DS | ACGCG1C†<br>C1GCGCA | 50.5±9.3 | 137.9±29.8 | 7.7±0.10 | 50.2 | 41.9±1.9 | 110.4±6.2 | 7.7±0.10 | 52.4 |
| DS | ACGCG1G<br>G1GCGCA | 57.5±7.6 | 155.1±23.3 | 9.41±0.39 | 58.5 | 50.7±8.3 | 134.2±25.6 | 9.11±0.52 | 59.5 |
| DS | ACGCG11<br>11GCGCA | 48.4±5.2 | 130.6±16.6 | 7.95±0.17 | 52.3 | 45.1±6.4 | 120.1±20.2 | 7.88±0.31 | 52.9 |
| DS | 1CGCGA1<br>1AGCGC1 | 50.8±4.7 | 136.9±14.8 | 8.35±0.17 | 54.2 | 45.5±6.7 | 120.2±21.3 | 8.26±0.35 | 55.6 |
| DS | GACGUC1<br>1CUGCAG | 71.2±6.4 | 197.9±19.5 | 9.86±0.35 | 56.4 | 64.9±6.8 | 178.3±21.1 | 9.59±0.32 | 56.9 |
| DS | CUGCAG1†<br>1GACGUC | 61.2±6.2 | 171.4±19.5 | 8.07±0.24 | 49.6 | 49.0±6.3 | 132.8±19.9 | 7.84±0.26 | 51.3 |
| DS | A1CGCGA<br>AGCGC1A | 58.5±9.2 | 162.6±29.3 | 8.11±0.14 | 50.5 | 56.1±5.7 | 154.7±18.0 | 8.12±0.18 | 51.1 |
| DS | C1CGCGA<br>AGCGC1C | 52.5±4.4 | 144.5±14.1 | 7.69±0.10 | 49.4 | 48.2±5.1 | 130.6±16.2 | 7.67±0.17 | 50.4 |
| DS | G1CGCGA<br>AGCGC1G | 55.0±6.1 | 152.0±19.5 | 7.90±0.10 | 50.1 | 53.2±4.6 | 146.0±14.7 | 7.89±0.13 | 50.5 |
| DS | 11CGCGA<br>AGCGC11 | 59.2±8.6 | 166.3±27.7 | 7.58±0.17 | 47.3 | 60.8±11.2 | 171.6±35.5 | 7.61±0.48 | 47.2 |
| DS | 1ACGCG1<br>1GCGCA1 | 50.6±10.1 | 139.0±31.7 | 7.49±0.31 | 48.5 | 51.1±23.3 | 140.7±72.7 | 7.50±2.20 | 48.4 |
| DS | 1CUGCAG†<br>GACGUC1 | 64.7±7.8 | 184.2±24.5 | 7.55±0.20 | 46.3 | 76.6±12.1 | 222.4±38.7 | 7.64±0.36 | 45.2 |
| DS | 1GACGUC<br>CUGCAG1 | 67.0±5.4 | 192.0±16.8 | 7.50±0.17 | 45.7 | 66.3±11.4 | 189.8±36.4 | 7.48±0.42 | 45.7 |

†These duplexes have a difference in $\Delta H°$ between the two fit methods that is greater than 15%. This can indicate non-two-state melting, but they were included in the analysis.
‡$T_M$ is calculated for strand concentration of 0.1 mM.

Table S8. Terminal mismatch systems studied by optical melting. The character "1" is used to represent 1mΨ. Site is either the Institute of Bioorganic Chemistry (IBCh) or DNA Software (DS).

| Site | Duplexes (5'-3') | Average of curve fits | | | | $T_M^{-1}$ vs log $C_T$ plots | | | |
|---|---|---|---|---|---|---|---|---|---|
| | | -ΔH° (kcal/mol) | -ΔS° (eu) | -ΔG°37 (kcal/mol) | $T_M$‡ (°C) | -ΔH° (kcal/mol) | -ΔS° (eu) | -ΔG°37 (kcal/mol) | $T_M$‡ (°C) |
| IBCh | A1GCGCAA<br>AACGCG1A | 60.3±1.3 | 159.4±3.9 | 10.89±0.12 | 66.2 | 64.4±4.3 | 171.6±12.7 | 11.18±0.32 | 66.0 |
| IBCh | UCAGUCAG1<br>AGUCAGUC1 | 83.1±3.6 | 230.1±10.9 | 11.75±0.23 | 57.7 | 78.1±4.0 | 215.3±12.0 | 11.45±0.24 | 57.3 |
| IBCh | AAUGCA1C<br>C1ACGUAA | 46.0±3.9 | 128.0±12.5 | 6.29±0.14 | 41.3 | 45.8±2.9 | 127.5±9.3 | 6.27±0.07 | 41.0 |
| IBCh | A1GCGCAC<br>CACGCG1A | 60.4±1.5 | 159.5±4.6 | 10.88±0.17 | 66.6 | 58.2±3.1 | 153.0±9.2 | 10.73±0.22 | 66.6 |
| IBCh | CAUGCA1A<br>A1ACGUAC | 49.4±0.9 | 136.7±2.5 | 7.04±0.13 | 45.6 | 53.8±3.4 | 150.4±10.9 | 7.12±0.09 | 45.8 |
| IBCh | 1GCGC1<br>1CGCG1 | 49.2±2.3 | 133.4±7.0 | 7.83±0.13 | 51.2 | 52.1±4.3 | 142.4±13.4 | 7.93±0.18 | 51.1 |
| IBCh | UCAGUCAG1<br>AGUCAGUCU | 80.7±3.1 | 221.4±9.3 | 12.05±0.23 | 59.7 | 81.6±3.0 | 223.9±9.1 | 12.13±0.21 | 60.0 |
| IBCh | UCAGUCAG1<br>AGUCAGUCC | 79.5±2.6 | 217.5±7.8 | 12.02±0.22 | 60.1 | 89.2±7.3 | 246.6±21.8 | 12.73±0.56 | 60.1 |
| IBCh | UAUGCAU1<br>1UACGUAU | 49.2±5.1 | 138.3±16.2 | 6.29±0.09 | 41.0 | 50.4±1.5 | 142.2±5.0 | 6.33±0.02 | 40.9 |
| IBCh | UAUGCA1U<br>U1ACGUAU | 50.4±1.7 | 140.9±5.4 | 6.67±0.06 | 43.4 | 55.1±1.0 | 156.0±3.2 | 6.70±0.01 | 43.0 |
| DS | AACGCG1A†<br>A1GCGCAA | 57.7±11.4 | 153.8±35.0 | 10.02±0.60 | 62.2 | 49.3±5.0 | 128.0±15.4 | 9.59±0.26 | 63.8 |
| DS | GACGCG1G†<br>G1GCGCAG | 64.8±7.0 | 175.4±21.5 | 10.39±0.39 | 61.3 | 54.0±4.2 | 142.3±12.8 | 9.84±0.22 | 63.0 |
| DS | 1ACGCG1G<br>G1GCGCA1 | 68.5±7.7 | 186.5±23.8 | 10.68±0.38 | 61.4 | 67.6±9.5 | 183.6±29.0 | 10.60±0.58 | 61.4 |
| DS | CACGCG1A<br>A1GCGCAC | 63.4±2.1 | 171.4±6.3 | 10.19±0.10 | 60.8 | 59.4±4.1 | 159.4±12.4 | 9.99±0.21 | 61.3 |
| DS | GACGCG1A<br>A1GCGCAG | 71.1±8.0 | 194.6±24.4 | 10.75±0.46 | 60.8 | 68.7±5.2 | 187.4±15.9 | 10.60±0.28 | 60.9 |
| DS | GACGCG11†<br>11GCGCAG | 66.4±3.7 | 181.1±11.4 | 10.23±0.22 | 59.8 | 53.8±7.3 | 141.5±22.2 | 9.64±0.43 | 63.7 |
| DS | AACGCG1G<br>G1GCGCAA | 65.0±9.1 | 175.5±27.9 | 10.56±0.47 | 62.2 | 63.2±3.9 | 170.2±12.0 | 10.44±0.21 | 62.3 |
| DS | CACGCG11<br>11GCGCAC | 56.4±2.7 | 153.2±8.4 | 8.90±0.12 | 55.8 | 53.9±6.4 | 145.4±19.9 | 8.83±0.29 | 56.3 |
| DS | 1ACGCG11<br>11GCGCA1 | 55.0±5.8 | 147.2±18.2 | 9.32±0.16 | 59.0 | 62.8±5.5 | 171.5±17.1 | 9.58±0.23 | 57.5 |
| DS | CACGCG1C<br>C1GCGCAC | 52.6±4.2 | 140.9±12.9 | 8.86±0.20 | 57.0 | 45.2±4.2 | 118.2±12.9 | 8.60±0.19 | 58.4 |
| DS | 1ACGCG1C†<br>C1GCGCA1 | 52.5±2.7 | 140.9±8.4 | 8.77±0.13 | 56.4 | 42.3±3.1 | 109.1±9.6 | 8.46±0.13 | 58.8 |
| DS | AACGCG1C†<br>C1GCGCAA | 52.6±1.2 | 141.1±3.7 | 8.84±0.10 | 56.9 | 45.2±2.5 | 117.8±7.7 | 8.61±0.09 | 58.6 |

| | | | | | | | | | |
|---|---|---|---|---|---|---|---|---|---|
| DS | 11CGCGAG†<br>GAGCGC11 | 61.4±8.0 | 166.1±24.4 | 9.94±0.45 | 60.1 | 51.4±3.4 | 135.2±10.4 | 9.42±0.17 | 61.4 |
| DS | G1CGCGAG<br>GAGCGC1G | 59.9±2.6 | 161.4±8.0 | 9.83±0.16 | 60.1 | 54.3±4.4 | 144.3±13.6 | 9.58±0.22 | 61.0 |
| DS | A1CGCGAA<br>AAGCGC1A | 60.4±5.8 | 162.8±17.8 | 9.93±0.27 | 60.5 | 60.2±4.5 | 162.2±13.9 | 9.88±0.21 | 60.3 |
| DS | C1CGCGAC†<br>CAGCGC1C | 61.5±7.4 | 168.6±23.3 | 9.22±0.25 | 55.9 | 48.0±6.6 | 126.3±20.7 | 8.85±0.32 | 59.0 |
| DS | C1CGCGAA<br>AAGCGC1C | 53.8±4.9 | 142.7±15.1 | 9.54±0.28 | 61.0 | 49.3±5.5 | 129.0±16.8 | 9.32±0.29 | 61.7 |
| DS | G1CGCGAA<br>AAGCGC1G | 62.4±4.2 | 169.2±13.0 | 9.92±0.22 | 59.6 | 55.4±3.3 | 147.8±10.0 | 9.59±0.15 | 60.6 |
| DS | A1CGCGAC<br>CAGCGC1A | 58.4±5.9 | 157.2±17.9 | 9.61±0.31 | 59.4 | 55.3±5.5 | 147.7±17.2 | 9.47±0.26 | 59.9 |
| DS | 11CGCGA1<br>1AGCGC11 | 62.9±4.6 | 170.9±14.1 | 9.86±0.27 | 59.1 | 55.6±4.6 | 148.7±14.1 | 9.53±0.22 | 60.1 |
| DS | A1CGCGAG†<br>GAGCGC1A | 66.3±5.5 | 180.5±16.6 | 10.28±0.34 | 60.1 | 56.0±6.3 | 149.0±19.4 | 9.76±0.35 | 61.4 |
| DS | G1CGCGA1<br>1AGCGC1G | 76.9±6.1 | 211.0±18.5 | 11.48±0.42 | 62.3 | 67.2±6.4 | 181.6±19.6 | 10.9±0.40 | 63.1 |
| DS | 11CGCGAC<br>CAGCGC11 | 58.2±5.8 | 158.2±17.7 | 9.18±0.32 | 56.9 | 52.6±5.2 | 140.8±16.2 | 8.95±0.24 | 57.6 |
| DS | C1CGCGA1<br>1AGCGC1C | 62.2±3.8 | 169.4±11.6 | 9.62±0.23 | 58.0 | 56.3±7.1 | 151.3±21.9 | 9.37±0.38 | 58.8 |
| DS | GCUGCAG1<br>1GACGUCG | 85.1±7.9 | 236.7±24.1 | 11.68±0.45 | 60.6 | 85.1±3.8 | 236.8±11.5 | 11.65±0.21 | 60.4 |
| DS | 1CUGCAGG<br>GGACGUC1 | 67.5±5.5 | 186.4±17.2 | 9.71±0.24 | 56.7 | 60.9±7.8 | 165.8±24.1 | 9.44±0.39 | 57.5 |
| DS | 1CUGCAG1†<br>1GACGUC1 | 64.1±2.9 | 176.6±8.9 | 9.34±0.22 | 55.8 | 49.5±7.0 | 130.9±21.8 | 8.92±0.34 | 58.7 |
| DS | CCUGCAG1<br>1GACGUCC | 68.0±5.6 | 189.0±17.6 | 9.39±0.16 | 54.9 | 66.4±6.9 | 184.0±21.5 | 9.36±0.29 | 55.2 |
| DS | 1CUGCAGC†<br>CGACGUC1 | 57.7±5.2 | 158.8±16.3 | 8.42±0.17 | 52.5 | 67.3±6.0 | 189.2±18.8 | 8.64±0.18 | 51.3 |
| DS | 1GACGUCG<br>GCUGCAG1 | 70.4±6.6 | 193.9±20.0 | 10.26±0.38 | 58.6 | 63.0±4.5 | 171.1±14.0 | 9.91±0.22 | 59.4 |
| DS | CGACGUC1<br>1CUGCAGC | 71.6±4.1 | 197.3±12.5 | 10.43±0.18 | 59.1 | 74.2±3.8 | 205.3±11.7 | 10.53±0.18 | 58.7 |
| DS | 1GACGUC1<br>1CUGCAG1 | 72.3±6.0 | 200.5±18.8 | 10.1±0.19 | 57.2 | 66.0±3.5 | 180.9±10.8 | 9.88±0.14 | 58.1 |
| DS | 1GACGUCC<br>CCUGCAG1 | 61.2±2.9 | 168.1±8.9 | 9.09±0.18 | 55.3 | 54.3±4.6 | 146.6±14.5 | 8.85±0.19 | 56.3 |
| DS | GGACGUC1<br>1CUGCAGG | 68.7±3.6 | 188.6±10.7 | 10.15±0.27 | 58.6 | 61.7±10.5 | 167.2±32.5 | 9.88±0.58 | 59.6 |

†These duplexes have a difference in ΔH° between the two fit methods that is greater than 15%. This can indicate non-two-state melting, but they were included in the analysis.

‡$T_M$ is calculated for strand concentration of 0.1 mM.

Table S9. Dangling end motif stability nearest neighbor parameters. Source indicates whether the parameter was measured in an experiment or estimated as explained in the text. The U analog parameter is from Turner 2004 parameters as tabulated by Zuber et al.[22]. The character "1" is used to represent 1mΨ.

| Sequence | Source | $\Delta G^\circ_{37}$ (kcal/mol) | U Analog $\Delta G^\circ_{37}$ (kcal/mol) | $\Delta\Delta G^\circ_{37}$ (kcal/mol) |
|---|---|---|---|---|
| 1A<br>U | Experiment | -0.53±0.20 | -0.20±0.14 | -0.33±0.24 |
| 1C<br>G | Experiment | -0.17±0.21 | -0.07±0.09 | -0.10±0.23 |
| 1G<br>C | Experiment | -0.38±0.21 | -0.15±0.12 | -0.23±0.24 |
| 1U<br>A | Estimate | -0.41±0.13 | -0.19±0.11 | -0.22±0.07 |
| 1G<br>U | Estimate | -0.42±0.16 | -0.20±0.14 | -0.22±0.07 |
| 1U<br>G | Estimate | -0.41±0.13 | -0.19±0.11 | -0.22±0.07 |
| AA<br>1 | Experiment | -0.48±0.22 | -0.30±0.13 | -0.18±0.25 |
| CA<br>1 | Experiment | -0.26±0.21 | -0.14±0.11 | -0.12±0.24 |
| GA<br>1 | Experiment | -0.37±0.21 | -0.20±0.14 | -0.17±0.25 |
| UA<br>1 | Estimate | -0.39±0.15 | -0.23±0.14 | -0.16±0.02 |
| 1A<br>1 | Experiment | -0.23±0.21 | -0.20±0.14 | -0.03±0.25 |
| A1<br>A | Estimate | -0.49±0.12 | -0.33±0.12 | -0.16±0.02 |
| C1<br>A | Estimate | -0.41±0.10 | -0.25±0.10 | -0.16±0.02 |
| G1<br>A | Estimate | -0.51±0.12 | -0.35±0.12 | -0.16±0.02 |
| U1<br>A | Estimate | -0.35±0.11 | -0.19±0.11 | -0.16±0.02 |
| 11<br>A | Experiment | -0.35±0.20 | -0.19±0.11 | -0.16±0.23 |
| A1<br>U | Experiment | -0.39±0.19 | -0.58±0.30 | 0.19±0.36 |
| C1<br>G | Experiment | -1.22±0.24 | -1.19±0.12 | -0.03±0.27 |
| G1<br>C | Experiment | -0.48±0.21 | -0.62±0.13 | 0.15±0.25 |

| | | | | |
|---|---|---|---|---|
| U1 A | Estimate | 0.01±0.18 | -0.09±0.17 | 0.10±0.07 |
| G1 U | Estimate | -0.48±0.31 | -0.58±0.30 | 0.10±0.07 |
| U1 G | Estimate | 0.01±0.18 | -0.09±0.17 | 0.10±0.07 |
| AA 1 | Experiment | -0.26±0.35 | -0.74±0.29 | 0.49±0.46 |
| AC 1 | Estimate | 0.00±0.48 | -0.49±0.15 | 0.49±0.46 |
| AG 1 | Estimate | -0.35±0.54 | -0.83±0.29 | 0.49±0.46 |
| AU 1 | Estimate | -0.09±0.55 | -0.58±0.30 | 0.49±0.46 |
| A1 1 | Experiment | -0.56±0.22 | -0.58±0.30 | 0.03±0.37 |
| 1A A | Experiment | -1.56±0.24 | -0.66±0.16 | -0.90±0.29 |
| 1C A | Experiment | -0.44±0.21 | -0.13±0.13 | -0.31±0.24 |
| 1G A | Experiment | -0.73±0.21 | -0.66±0.13 | -0.07±0.24 |
| 1U A | Estimate | -0.52±0.17 | -0.09±0.17 | -0.43±0.02 |
| 11 A | Experiment | -0.54±0.21 | -0.09±0.17 | -0.45±0.27 |

Table S10. Terminal mismatch motif stability nearest neighbor parameters. Source indicates whether the parameter was measured in an experiment or estimated as explained in the text. The U analog parameter is from Turner 2004 parameters as tabulated by Zuber et al.[22]. The character "1" is used to represent 1mΨ. $\Delta\Delta G^{\circ}_{37}$ is the stability change of the mismatch with 1mΨ relative to that with U.

| Mismatch | Source | $\Delta G^{\circ}_{37}$ (kcal/mol) | U Analog $\Delta G^{\circ}_{37}$ (kcal/mol) | $\Delta\Delta G^{\circ}_{37}$ (kcal/mol) |
|---|---|---|---|---|
| AC<br>U1 | Estimate | -0.94±0.34 | -0.71±0.21 | -0.23±0.28 |
| AU<br>U1 | Estimate | -1.13±0.47 | -0.80±0.33 | -0.33±0.33 |
| A1<br>UC | Estimate | -1.32±0.56 | -0.72±0.32 | -0.60±0.46 |
| A1<br>UU | Estimate | -1.25±0.66 | -0.80±0.33 | -0.45±0.57 |
| A1<br>U1 | Estimate | -1.13±0.47 | -0.80±0.33 | -0.33±0.33 |
| CC<br>G1 | Experiment | -0.85±0.23 | -0.76±0.42 | -0.09±0.48 |
| CU<br>G1 | Estimate | -1.52±0.62 | -1.19±0.52 | -0.33±0.33 |
| C1<br>GC | Experiment | -1.69±0.25 | -1.36±0.53 | -0.33±0.59 |
| C1<br>GU | Estimate | -2.29±0.84 | -1.19±0.52 | -1.10±0.66 |
| C1<br>G1 | Experiment | -1.51±0.15 | -1.19±0.52 | -0.32±0.54 |
| GC<br>C1 | Experiment | -0.88±0.22 | -0.50±0.16 | -0.38±0.27 |
| GU<br>C1 | Estimate | -1.10±0.37 | -0.77±0.18 | -0.33±0.33 |
| G1<br>CC | Experiment | -1.85±0.69 | -0.98±0.18 | -0.87±0.72 |
| G1<br>CU | Experiment | -1.87±0.64 | -0.77±0.18 | -1.10±0.66 |
| G1<br>C1 | Experiment | -1.10±0.33 | -0.77±0.18 | -0.33±0.37 |
| UC<br>A1 | Estimate | -0.69±0.61 | -0.46±0.55 | -0.23±0.28 |
| UU<br>A1 | Estimate | -0.84±0.62 | -0.51±0.53 | -0.33±0.33 |
| U1<br>AC | Estimate | -1.19±0.81 | -0.59±0.67 | -0.6±0.46 |
| U1<br>AU | Experiment | -0.96±0.21 | -0.51±0.53 | -0.45±0.57 |

| | | | | |
|---|---|---|---|---|
| U1<br>A1 | Estimate | -0.84±0.62 | -0.51±0.53 | -0.33±0.33 |
| GC<br>U1 | Estimate | -0.94±0.34 | -0.71±0.21 | -0.23±0.28 |
| GU<br>U1 | Estimate | -1.13±0.47 | -0.80±0.33 | -0.33±0.33 |
| G1<br>UC | Estimate | -1.32±0.56 | -0.72±0.32 | -0.60±0.46 |
| G1<br>UU | Estimate | -1.25±0.66 | -0.80±0.33 | -0.45±0.57 |
| G1<br>U1 | Estimate | -1.13±0.47 | -0.80±0.33 | -0.33±0.33 |
| UC<br>G1 | Estimate | -0.69±0.61 | -0.76±0.42 | -0.09±0.48 |
| UU<br>G1 | Estimate | -0.84±0.62 | -1.19±0.52 | -0.33±0.33 |
| U1<br>GC | Estimate | -1.19±0.81 | -1.36±0.53 | -0.33±0.59 |
| U1<br>GU | Estimate | -0.96±0.21 | -1.19±0.52 | -1.10±0.66 |
| U1<br>G1 | Estimate | -0.84±0.62 | -1.19±0.52 | -0.32±0.54 |
| AA<br>1A | Experiment | -1.02±0.22 | -0.75±0.42 | -0.27±0.47 |
| AA<br>1C | Experiment | -1.09±0.23 | -0.88±0.31 | -0.21±0.39 |
| AA<br>1G | Experiment | -1.22±0.24 | -0.78±0.18 | -0.45±0.3 |
| AC<br>1A | Experiment | -0.80±0.22 | -0.62±0.41 | -0.18±0.46 |
| AC<br>1C | Experiment | -0.85±0.23 | -0.63±0.19 | -0.22±0.29 |
| AC<br>1U | Estimate | -0.67±0.36 | -0.71±0.21 | 0.04±0.41 |
| AC<br>11 | Experiment | -0.90±0.23 | -0.71±0.21 | -0.19±0.31 |
| AG<br>1A | Experiment | -1.31±0.24 | -0.80±0.29 | -0.51±0.38 |
| AG<br>1G | Experiment | -1.22±0.24 | -0.78±0.26 | -0.44±0.35 |
| AU<br>1C | Estimate | -0.51±0.52 | -0.72±0.32 | 0.21±0.61 |
| AU<br>1U | Estimate | -0.86±0.41 | -0.8±0.33 | -0.06±0.53 |
| AU<br>11 | Estimate | -0.33±0.59 | -0.80±0.33 | 0.48±0.67 |

| | | | | |
|---|---|---|---|---|
| A1<br>1C | Experiment | -1.11±0.24 | -0.72±0.32 | -0.39±0.40 |
| A1<br>1U | Estimate | -1.31±0.70 | -0.80±0.33 | -0.51±0.77 |
| A1<br>11 | Experiment | -1.19±0.24 | -0.80±0.33 | -0.39±0.41 |
| 1A<br>AA | Experiment | -1.40±0.24 | -0.94±0.52 | -0.46±0.57 |
| 1A<br>AC | Experiment | -1.60±0.24 | -0.76±0.53 | -0.84±0.58 |
| 1A<br>AG | Experiment | -1.90±0.25 | -1.10±0.42 | -0.80±0.49 |
| 1C<br>AA | Experiment | -0.91±0.22 | -0.72±0.39 | -0.19±0.45 |
| 1C<br>AC | Experiment | -0.90±0.22 | -0.59±0.42 | -0.31±0.47 |
| 1C<br>AU | Estimate | -0.60±0.35 | -0.46±0.55 | -0.14±0.65 |
| 1C<br>A1 | Experiment | -0.83±0.22 | -0.46±0.55 | -0.37±0.59 |
| 1G<br>AA | Experiment | -1.82±0.25 | -1.11±0.58 | -0.71±0.63 |
| 1G<br>AG | Experiment | -1.52±0.24 | -1.15±0.59 | -0.37±0.64 |
| 1U<br>AC | Estimate | -0.41±0.51 | -0.59±0.67 | 0.18±0.84 |
| 1U<br>AU | Experiment | -0.14±0.34 | -0.51±0.53 | 0.37±0.63 |
| 1U<br>A1 | Estimate | -0.87±0.26 | -0.51±0.53 | -0.36±0.59 |
| 11<br>AC | Experiment | -1.02±0.22 | -0.59±0.67 | -0.43±0.71 |
| 11<br>AU | Estimate | -0.59±0.66 | -0.51±0.53 | -0.08±0.85 |
| 11<br>A1 | Experiment | -1.39±0.23 | -0.51±0.53 | -0.88±0.58 |
| GA<br>1A | Estimate | -1.02±0.22 | -0.25±0.56 | -0.77±0.60 |
| GA<br>1C | Estimate | -1.09±0.23 | -0.88±0.31 | -0.21±0.39 |
| GA<br>1G | Estimate | -1.22±0.24 | -0.78±0.18 | -0.45±0.30 |
| GC<br>1A | Estimate | -0.80±0.22 | -0.62±0.41 | -0.18±0.46 |
| GC<br>1C | Estimate | -0.85±0.23 | -0.63±0.19 | -0.22±0.29 |

| | | | | |
|---|---|---|---|---|
| GC<br>1U | Estimate | -0.67±0.36 | -0.71±0.21 | 0.04±0.41 |
| GC<br>11 | Estimate | -0.90±0.23 | -0.71±0.21 | -0.19±0.31 |
| GG<br>1A | Estimate | -1.31±0.24 | -0.58±0.42 | -0.73±0.48 |
| GG<br>1G | Estimate | -1.22±0.24 | -0.78±0.26 | -0.44±0.35 |
| GU<br>1C | Estimate | -0.51±0.52 | -0.72±0.32 | 0.21±0.61 |
| GU<br>1U | Estimate | -0.86±0.41 | -0.57±0.54 | -0.29±0.68 |
| GU<br>11 | Estimate | -0.33±0.59 | -0.57±0.54 | 0.25±0.80 |
| G1<br>UC | Estimate | -1.11±0.24 | -0.72±0.32 | -0.39±0.40 |
| G1<br>1U | Estimate | -1.31±0.70 | -0.57±0.54 | -0.74±0.88 |
| G1<br>11 | Estimate | -1.19±0.24 | -0.57±0.54 | -0.62±0.59 |
| 1A<br>GA | Estimate | -1.40±0.24 | -0.94±0.52 | -0.46±0.57 |
| 1A<br>GC | Estimate | -1.60±0.24 | -0.76±0.53 | -0.84±0.58 |
| 1A<br>GG | Estimate | -1.90±0.25 | -1.10±0.42 | -0.80±0.49 |
| 1C<br>GA | Estimate | -0.91±0.22 | -0.72±0.39 | -0.19±0.45 |
| 1C<br>GC | Estimate | -0.90±0.22 | -0.59±0.42 | -0.31±0.47 |
| 1C<br>GU | Estimate | -0.60±0.35 | -0.46±0.55 | -0.14±0.65 |
| 1C<br>G1 | Estimate | -0.83±0.22 | -0.46±0.55 | -0.37±0.59 |
| 1G<br>GA | Estimate | -1.82±0.25 | -0.48±0.56 | -1.34±0.61 |
| 1G<br>GG | Estimate | -1.52±0.24 | -0.81±0.55 | -0.71±0.60 |
| 1U<br>GC | Estimate | -0.41±0.51 | -0.59±0.67 | 0.18±0.84 |
| 1U<br>GU | Estimate | -0.14±0.34 | -0.40±0.22 | 0.26±0.40 |
| 1U<br>G1 | Estimate | -0.87±0.26 | -0.40±0.22 | -0.47±0.34 |
| 11<br>GC | Estimate | -1.02±0.22 | -0.59±0.67 | -0.43±0.71 |

| | | | | |
|---|---|---|---|---|
| 11<br>GU | Estimate | -0.59±0.66 | -0.40±0.22 | -0.19±0.70 |
| 11<br>G1 | Estimate | -1.39±0.23 | -0.40±0.22 | -0.99±0.32 |

Table S11. Hairpin loops studied by optical melting, where unpaired nucleotides are underlined. The character "1" is used to represent 1mΨ. All hairpin loops were studied at the Institute of Bioorganic Chemistry.

| Hairpin Sequence | $-\Delta H^{\circ}$ (kcal/mol) | $-\Delta S^{\circ}$ (eu) | $-\Delta G^{\circ}_{37}$ (kcal/mol) | $T_M$ (˚C) |
|---|---|---|---|---|
| CGUGUUCGAUCCACG | 28.1±5.2 | 85.4±15.6 | 1.56±0.38 | 55.3 |
| CGUGU1CGAUCCACG | 38.2±5.1 | 116.3±15.3 | 2.13±0.36 | 55.3 |
| CAGACUGAAGAUCUG | 16.1±3.5 | 51.0±10.7 | 0.24±0.13 | 41.8 |
| CAGACUGAAGA1CUG | 43.1±5.0 | 129.4±15.3 | 2.94±0.24 | 59.7 |
| GGAUUAAUUUCC | 40.7±2.0 | 126.3±6.2 | 1.58±0.14 | 49.5 |
| GGAUUAAUU1CC | 41.2±2.3 | 126.9±7.2 | 1.83±0.07 | 51.4 |
| GGA1UAAUUUCC | 40.9±3.3 | 125.4±9.9 | 1.97±0.24 | 52.7 |
| GGA1UAAUU1CC | 40.2±1.7 | 122.8±4.9 | 2.12±0.16 | 54.3 |
| GGACUUCGGUCC | 42.2±5.8 | 120.7±16.7 | 4.78±0.61 | 76.6 |
| GGAC1UCGGUCC | 52.2±3.3 | 148.6±9.9 | 6.14±0.23 | 78.3 |
| GGACU1CGGUCC | 51.7±2.4 | 149.6±7.0 | 5.26±0.23 | 72.2 |
| GGACUUCGG1CC | 53.7±1.7 | 152.0±5.3 | 6.52±0.08 | 79.9 |

Table S12. Hairpin loop stabilities. The character "1" is used to represent 1mΨ.

| Sequence | $\Delta G^{\circ}_{37\ loop}$ Measured (kcal/mol) | $\Delta G^{\circ}_{37\ loop}$ Estimated† (kcal/mol) | $\Delta\Delta G^{\circ}_{37}$ (kcal/mol) |
|---|---|---|---|
| CGUGUUCGAUCCACG | 5.15±0.14 | 5.21±0.64 | -0.06±0.66 |
| CGUGU1CGAUCCACG | 4.58±0.15 | 5.21±0.64 | -0.63±0.66 |
| CAGACUGAAGAUCUG | 5.85±0.12 | 5.57±0.74 | 0.28±0.75 |
| CAGACUGAAGA1CUG | 3.78±0.29 | 5.39±0.66 | -1.61±0.72 |
| GGAUUAAUUUCC | 3.58±0.12 | 3.73±0.98 | -0.15±0.99 |
| GGAUUAAUU1CC | 3.96±0.27 | 3.67±1.01 | 0.29±1.04 |
| GGA1UAAUUUCC | 3.19±0.13 | 3.29±1.13 | -0.10±1.14 |
| GGA1UAAUU1CC | 3.67±0.27 | 3.22±1.16 | 0.45±1.19 |
| GGACUUCGGUCC | 3.07±0.22 | 2.21±0.37 | 0.86±0.43 |
| GGAC1UCGGUCC | 1.71±0.27 | 2.21±0.37 | -0.5±0.46 |
| GGACU1CGGUCC | 2.59±0.24 | 2.21±0.37 | 0.38±0.44 |
| GGACUUCGG1CC | 2.07±0.44 | 2.21±0.37 | -0.14±0.57 |

†Hairpin loop stabilities are estimated using the Turner 2004 nearest neighbor rules as modified for 1mΨ. For the UUCG tetraloops (including variations 1mΨ-UCG and U-1mΨ-CG, the values are from a lookup table.[15] For the other hairpin loops studied here, the functional form is:

$$\Delta G^{\circ}_{37\ loop} = \Delta G^{\circ}_{37\ initiation}(n) + \Delta G^{\circ}_{37}(\text{terminal mismatch}) + \Delta G^{\circ}_{37\ initiation}(\text{UU first mismatch})$$

where n is the number of unpaired nucleotides, $\Delta G^{\circ}_{37}$(terminal mismatch) is the terminal mismatch stability based on the sequence of the closing pair and first mismatch (Table S10 for motifs with 1mΨ), and $\Delta G^{\circ}_{37}$(UU first mismatch) is an additional stability that applies to U-U, U-1mΨ, 1mΨ-U, and 1mΨ -1mΨ first mismatches.

Table S13. Internal loop models studied by optical melting. The character "1" is used to represent 1mΨ. Site is either the Institute of Bioorganic Chemistry (IBCh) or DNA Software (DS).

| | | Average of curve fits | | | | $T_M^{-1}$ vs log $C_T$ plots | | | |
|---|---|---|---|---|---|---|---|---|---|
| Sequence | Site: | -ΔH˚ (kcal/mol) | -ΔS˚ (eu) | $-\Delta G^\circ_{37}$ (kcal/mol) | $T_M$† (˚C) | -ΔH˚ (kcal/mol) | -ΔS˚ (eu) | $-\Delta G^\circ_{37}$ (kcal/mol) | $T_M$† (˚C) |
| C1GC1GG<br>GG1CG1C | IBCh | 41.2±2.1 | 113.9±6.7 | 5.88±0.08 | 38.5 | 45.1±2.9 | 126.9±9.5 | 5.86±0.06 | 37.5 |
| CAUG11ACUAC<br>GUACAC1GAUG | DS | 92.0±7.2 | 263.7±22.8 | 10.21±0.10 | 49.9 | 95.1±4.8 | 273.5±15.2 | 10.27±0.11 | 49.7 |
| CAUGA1GCUAC<br>GUAC1CCGAUG | DS | 90.7±1.3 | 255.3±4.0 | 11.49±0.06 | 54.9 | 81.0±3.6 | 225.2±11.3 | 11.18±0.12 | 55.9 |
| CAUGACGCUAC<br>GUAC11CGAUG | DS | 88.7±2.1 | 249.7±6.5 | 11.22±0.08 | 54.3 | 86.0±1.5 | 241.3±4.8 | 11.13±0.05 | 54.5 |
| CCAAAGCA<br>AGG1G1CG | DS | 86.6±10.7 | 250.7±33.8 | 8.82±0.21 | 45.4 | 63.9±5.4 | 178.3±17.1 | 8.59±0.10 | 47.3 |
| CCACAGCA<br>AGG1A1CG | DS | 77.4±2.8 | 222.4±8.9 | 8.40±0.14 | 44.7 | 63.1±5.8 | 176.5±18.5 | 8.35±0.14 | 46.2 |
| CG1C1CAUGA1ACG<br>GCA1AGUAC1C1GC | DS | 69.5±11.6 | 190.4±35.9 | 10.46±0.49 | 59.9 | 68.7±6.3 | 188.0±19.4 | 10.36±0.29 | 59.7 |
| CGACCAUAUG11CG<br>GC11GUAUACCAGC | DS | 69.3±8.9 | 192.9±27.7 | 9.50±0.27 | 55.1 | 67.0±3.8 | 185.6±12.0 | 9.39±0.12 | 55.2 |
| CGC1GCG<br>GCG1CGC | IBCh | 66.2±6.5 | 184.7±20.6 | 8.92±0.19 | 53.0 | 68.7±6.4 | 192.7±19.8 | 9.00±0.26 | 52.2 |
| CUCCACAUG11GAG<br>GAG11GUACACCUC | DS | 95.1±7.6 | 268.9±23.5 | 11.70±0.29 | 58.0 | 109.6±4.1 | 313.6±12.7 | 12.34±0.19 | 57.1 |
| CUUG1CAAG<br>GAAC1GUUC | DS | 75.8±3.6 | 218.0±11.4 | 8.17±0.13 | 47.5 | 65.1±6.1 | 184.0±19.4 | 8.04±0.15 | 48.7 |
| GAACGC1GUCC<br>CUUGC1ACAGG | DS | 104.5±7.5 | 293.8±23.0 | 13.43±0.36 | 58.9 | 102.4±5.9 | 287.3±18.2 | 13.30±0.30 | 59.0 |
| GAC11AGUC<br>CUGA11CAG | DS | 66.9±4.8 | 188.3±15.1 | 8.52±0.17 | 50.8 | 60.4±2.9 | 167.6±9.3 | 8.38±0.06 | 51.5 |
| GAC1AAGUC<br>CUGAA1CAG | DS | 70.3±4.6 | 203.8±14.7 | 7.03±0.08 | 43.1 | 70.8±7.7 | 205.6±24.8 | 7.03±0.14 | 43.0 |
| GAC1AAGUC<br>CUGAA1CAG | DS | 69.3±3.9 | 200.4±12.3 | 7.17±0.09 | 43.8 | 80.1±6.7 | 235.0±21.4 | 7.19±0.09 | 43.0 |
| GAC1CACGUGCGUC<br>CUGCGUGCAC1GAC | DS | 125.8±7.7 | 354.0±23.1 | 16.01±0.48 | 64.8 | 141.8±11.3 | 402.2±33.8 | 17.09±0.78 | 64.1 |
| GAC1CAGUC<br>CUGAC1CAG | DS | 72.0±1.5 | 210.9±4.5 | 6.62±0.10 | 41.1 | 83.8±10.2 | 249.0±32.9 | 6.53±0.21 | 40.2 |
| GAC1GAGUC<br>CUGAG1CAG | DS | 80.2±6.3 | 232.3±19.8 | 8.16±0.16 | 46.9 | 81.2±8.0 | 235.6±25.4 | 8.16±0.17 | 46.8 |
| GACA11GUC<br>CUG11ACAG | DS | 70.9±4.6 | 200.5±14.3 | 8.7±0.19 | 50.8 | 55.3±0.7 | 151.4±2.2 | 8.38±0.01 | 52.9 |
| GACAA1GUC<br>CUG1AACAG | DS | 72.3±4.2 | 207.5±13.3 | 7.91±0.08 | 46.9 | 73.7±3.3 | 212.0±10.7 | 7.91±0.04 | 46.7 |
| GACAC1GUC<br>CUG1CACAG | DS | 71.6±13.6 | 209.9±44.3 | 6.84±0.15 | 40.7 | 75.5±10.3 | 221.6±33.4 | 6.77±0.26 | 41.5 |
| GACAG1GUC<br>CUG1GACAG | DS | 71.6±3.5 | 201.9±10.8 | 8.98±0.15 | 52.0 | 66.3±6.5 | 185.4±20.5 | 8.83±0.21 | 52.5 |

| | | | | | | | | | |
|---|---|---|---|---|---|---|---|---|---|
| GACC1CCUGUG<br>CUGGA1GACAC | DS | 106.0±5.3 | 296.0±16.1 | 14.21±0.28 | 61.2 | 119.0±7.4 | 335.4±22.6 | 14.92±0.42 | 60.5 |
| GAUC1GAUC<br>CUAG1CUAG | DS | 77.4±4.3 | 222.1±13.6 | 8.47±0.15 | 48.6 | 67.2±3.8 | 189.9±12.1 | 8.30±0.08 | 49.6 |
| GAUCA11GUAC<br>CUAG1CACAUG | DS | 89.8±9.8 | 257.0±31.3 | 10.10±0.12 | 49.9 | 92.3±6.7 | 264.9±21.1 | 10.13±0.16 | 49.6 |
| GC111CCA<br>ACGACAGG | DS | 83.1±7.2 | 240.7±23.1 | 8.44±0.11 | 44.3 | 68.5±4.9 | 193.9±15.6 | 8.39±0.07 | 45.6 |
| GC1A1CCA<br>ACGAGAGG | DS | 67.2±7.1 | 185.7±22.5 | 9.61±0.13 | 51.9 | 67.3±17.5 | 185.9±54.8 | 9.62±0.90 | 51.9 |
| GC1C1CCA<br>ACGAAAGG | DS | 69.6±7.0 | 196.2±22.2 | 8.75±0.16 | 47.2 | 56.4±4.8 | 154.0±15.1 | 8.61±0.13 | 48.9 |
| GCA1UCG<br>CGUUAGC | IBCh | 62.9±2.3 | 182.5±7.6 | 6.25±0.11 | 35.9 | 65.7±2.4 | 191.9±7.8 | 6.19±0.05 | 35.4 |
| GCGU1UCGC<br>CGCU1UGCG | IBCh | 65.2±4.6 | 187.8±15.1 | 6.97±0.09 | 43.2 | 72.5±2.2 | 211.4±7.1 | 6.99±0.02 | 42.5 |
| UCAG1CAGU<br>AGUC1GUCA | IBCh | 76.8±5.0 | 214.8±15.4 | 10.24±0.27 | 52.5 | 69.6±2.8 | 192.4±8.7 | 9.95±0.10 | 52.9 |
| CGAGC1CUCG<br>GCUC1CGAGC | DS | 82.6±6.6 | 225.7±20.0 | 12.58±0.42 | 65.3 | 84.8±7.6 | 232.3±22.9 | 12.70±0.51 | 65.0 |
| GAAUGAAG<br>CUU1G1UC | IBCh | 67.6±1.1 | 203.5±3.5 | 4.41±0.03 | 27.9 | 71.1±1.7 | 215.2±5.5 | 4.39±0.06 | 27.8 |
| GCA11UGC<br>CGU11ACG | IBCh | 66.6±4.2 | 190.8±12.9 | 7.43±0.21 | 45.5 | 68.1±3.3 | 195.5±10.2 | 7.42±0.06 | 45.4 |
| GCA1UUGC<br>CGUU1ACG | IBCh | 68.2±3.8 | 195.7±11.9 | 7.48±0.27 | 45.5 | 71.2±7.1 | 205.9±23.0 | 7.46±0.19 | 44.4 |
| GCAC1CGUGC<br>CGUGC1CACG | DS | 47.3±13.0 | 125.5±40.9 | 8.40±0.26 | 55.9 | 43.7±4.0 | 114.3±12.6 | 8.22±0.13 | 56.2 |
| GCACC1GUGC<br>CGUG1CCACG | DS | 75.1±8.7 | 215.4±27.9 | 8.27±0.10 | 48.1 | 74.3±10.7 | 212.7±33.8 | 8.29±0.34 | 48.3 |
| GCAG11CUGG<br>CGUCCCGACC | DS | 81.8±8.3 | 235.3±26.3 | 8.80±0.23 | 45.9 | 77.7±20.0 | 222.1±62.9 | 8.77±1.03 | 46.2 |
| GCAU11GC<br>CG11UACG | IBCh | 55.3±4.3 | 161.8±14.1 | 5.09±0.16 | 33.9 | 62.0±2.5 | 184.0±8.2 | 4.94±0.05 | 33.3 |
| GCAUU1GC<br>CG1UUACG | IBCh | 61.5±3.5 | 181.0±11.7 | 5.40±0.22 | 35.4 | 66.0±3.7 | 195.9±12.0 | 5.29±0.08 | 35.0 |
| GCUGG1GC<br>CG1GGUCG | IBCh | 64.7±5.8 | 182.0±18.1 | 8.24±0.26 | 49.9 | 60.7±4.8 | 169.4±15.3 | 8.11±0.15 | 50.2 |
| GGAC11CUGG<br>CCUGCCGACC | DS | 89.2±3.5 | 259.7±11.0 | 8.66±0.09 | 44.6 | 79.4±6.3 | 228.0±20.2 | 8.62±0.11 | 45.4 |
| GGAC11GUCC<br>CCUG11CAGG | DS | 93.2±10.1 | 259.6±30.7 | 12.63±0.56 | 62.0 | 91.2±7.5 | 253.7±22.7 | 12.50±0.42 | 62.1 |
| GGAG11CUCC<br>CCUC11GAGG | DS | 97.6±4.7 | 272.9±14.2 | 12.94±0.32 | 62.0 | 102.5±10.0 | 287.8±30.3 | 13.20±0.59 | 61.6 |
| GCG1U1CGC<br>CGC1U1GCG | IBCh | 71.9±5.3 | 203.4±16.6 | 8.85±0.19 | 51.2 | 74.2±3.7 | 210.5±11.6 | 8.90±0.14 | 51.1 |
| GCG1UUCGC<br>CGCUU1GCG | IBCh | 76.4±9.4 | 216.4±29.3 | 9.33±0.33 | 52.4 | 86.2±3.6 | 247.0±11.2 | 9.62±0.14 | 51.8 |

| | | | | | | | | | |
|---|---|---|---|---|---|---|---|---|---|
| CAUG1CACUAC<br>GUACA11GAUG | DS | 89.4±7.2 | 254.8±22.8 | 10.34±0.11 | 50.8 | 93.7±7.9 | 268.5±24.7 | 10.45±0.22 | 50.5 |

†$T_M$ is calculated for strand concentration of 0.1 mM.

Table S14. Internal loop stabilities. The character “1” is used to represent 1mΨ. Site is either the Institute of Bioorganic Chemistry (IBCh) or DNA Software (DS). $\Delta G^\circ_{37\ loop}$ is the stability of loop as shown, including 1mΨ. $\Delta G^\circ_{37\ loop}$ (U analog) is the stability of loop with all 1mΨ replaced with U. $\Delta G^\circ_{37\ loop}$ (Predicted) is the estimated stability of the loop using the nearest neighbor parameters derived in this work. $\Delta\Delta G^\circ_{37}$ is the difference in loop stability between the predicted and measured values.

| Sequence | Site | $\Delta G^\circ_{37\ loop}$ (kcal/mol) | U analog $\Delta G^\circ_{37\ loop}$ (kcal/mol) | $\Delta G^\circ_{37\ loop}$ (Predicted) (kcal/mol) | $\Delta\Delta G^\circ_{37}$ (Predicted-Measured) (kcal/mol) |
|---|---|---|---|---|---|
| C1GC1GG<br>GG1CG1C | IBCh | -2.48±1.03 | 0.94±1.02 | 0.94±1.02 | 3.42±1.45 |
| CAUG11ACUAC<br>GUACAC1GAUG | DS | 2.21±0.70 | 0.94±1.02 | 2.34±1.04 | 0.13±1.25 |
| CAUG1CACUAC<br>GUACA11GAUG | DS | 2.03±0.70 | 0.94±1.02 | 2.34±1.04 | 0.31±1.26 |
| CAUGA1GCUAC<br>GUAC1CCGAUG | DS | 1.65±0.60 | 0.63±0.62 | 1.33±0.62 | -0.32±0.87 |
| CAUGACGCUAC<br>GUAC11CGAUG | DS | 1.7±0.60 | 0.63±0.62 | 1.33±0.62 | -0.37±0.86 |
| CCAAAGCA<br>AGG1G1CG | DS | 1.68±0.63 | 0.94±1.02 | 2.34±1.04 | 0.66±1.22 |
| CCACAGCA<br>AGG1A1CG | DS | 1.92±0.62 | 0.94±1.02 | 2.34±1.04 | 0.42±1.21 |
| CG1C1CAUGA1ACG<br>GCA1AGUAC1C1GC | DS | 2.91±0.44 | 0.94±1.02 | 3.73±1.09 | 0.82±1.18 |
| CGACCAUAUG11CG<br>GC11GUAUACCAGC | DS | 1.81±0.36 | 0.63±0.62 | 2.03±0.65 | 0.22±0.74 |
| CGC1GCG<br>GCG1CGC | IBCh | -1.96±0.49 | 0.42±0.40 | -1.44±0.47 | 0.52±0.68 |
| CUCCACAUG11GAG<br>GAG11GUACACCUC | DS | 1.46±0.38 | 0.63±0.62 | 2.03±0.65 | 0.57±0.75 |
| CUUG1CAAG<br>GAAC1GUUC | DS | -2.32±0.44 | -0.7±0.41 | -2.56±0.48 | -0.24±0.65 |
| GAACGC1GUCC<br>CUUGC1ACAGG | DS | 0.92±0.66 | 0.63±0.62 | 1.33±0.62 | 0.41±0.91 |
| GAC11AGUC<br>CUGA11CAG | DS | 0.72±0.65 | 1.80±0.48 | 1.34±0.57 | 0.62±0.87 |
| GAC1AAGUC<br>CUGAA1CAG | DS | 2.07±0.62 | 0.94±1.02 | 2.34±1.04 | 0.27±1.21 |
| GAC1AAGUC<br>CUGAA1CAG | DS | 1.91±0.62 | 0.94±1.02 | 2.34±1.04 | 0.43±1.21 |
| GAC1CACGUGCGUC<br>CUGCGUGCAC1CAG | DS | -0.69±0.39 | -0.03±0.47 | -0.03±0.47 | 0.66±0.61 |
| GAC1CAGUC<br>CUGAC1CAG | DS | 2.57±0.61 | 0.94±1.02 | 2.34±1.04 | -0.23±1.21 |
| GAC1GAGUC<br>CUGAG1CAG | DS | 0.94±0.64 | -1.60±1.10 | -0.20±1.12 | -1.14±1.29 |

| | | | | | |
|---|---|---|---|---|---|
| GACA11GUC<br>CUG11ACAG | DS | 1.44±0.66 | 0.38±1.32 | -0.08±1.36 | -1.52±1.51 |
| GACAA1GUC<br>CUG1AACAG | DS | 1.91±0.65 | 0.94±1.02 | 2.34±1.04 | 0.43±1.23 |
| GACAC1GUC<br>CUG1CACAG | DS | 3.05±0.63 | 0.94±1.02 | 2.34±1.04 | -0.71±1.22 |
| GACAG1GUC<br>CUG1GACAG | DS | 0.99±0.67 | -1.60±1.10 | -0.20±1.12 | -1.19±1.31 |
| GACC1CCUGUG<br>CUGGA1GACAC | DS | -0.40±0.71 | 0.63±0.62 | 1.33±0.62 | 1.73±0.95 |
| GAUC1GAUC<br>CUAG1CUAG | DS | -1.22±0.49 | 0.41±0.40 | -1.45±0.47 | -0.23±0.68 |
| GAUCA11GUAC<br>CUAG1CACAUG | DS | 2.55±0.71 | 0.94±1.02 | 2.34±1.04 | -0.21±1.26 |
| GC111CCA<br>ACGACAGG | DS | 2.41±0.79 | 0.94±1.02 | 2.34±1.04 | -0.07±1.31 |
| GC1A1CCA<br>ACGAGAGG | DS | 1.18±0.82 | 0.94±1.02 | 2.34±1.04 | 1.16±1.32 |
| GC1C1CCA<br>ACGAAAGG | DS | 2.19±0.80 | 0.94±1.02 | 2.34±1.04 | 0.15±1.31 |
| GCA1UCG<br>CGUUAGC | IBCh | -0.94±0.37 | 0.38±1.32 | -1.03±1.33 | -0.09±1.38 |
| UCAG1CAGU<br>AGUC1GUCA | IBCh | -1.97±0.5 | -0.7±0.41 | -2.56±0.43 | -0.59±0.66 |
| CGAGC1CUCG<br>GCUC1CGAGC | DS | -3.64±0.61 | 1.05±0.30 | 1.05±0.30 | 4.69±0.68 |
| GAAUGAAG<br>CUU1G1UC | IBCh | -2.45±0.41 | 1.48±0.11 | 2.29±0.15 | 4.74±0.44 |
| GCA11UGC<br>CGU11ACG | IBCh | -1.78±0.43 | 0.60±0.12 | -1.26±0.28 | 0.52±0.51 |
| GCA1UUGC<br>CGUU1ACG | IBCh | -1.82±0.43 | 0.60±0.12 | -2.22±0.32 | -0.4±0.54 |
| GCAC1CGUGC<br>CGUGC1CACG | DS | 2.80±0.46 | 1.36±0.24 | 1.36±0.24 | -1.44±0.52 |
| GCACC1GUGC<br>CGUC1CCACG | DS | 2.73±0.47 | 1.29±0.20 | 1.29±0.20 | -1.44±0.51 |
| GCAG11CUGG<br>CGUCCCGACC | DS | 2.2±0.47 | 1.01±0.20 | 1.01±0.20 | -1.19±0.51 |
| GCAU11GC<br>CG11UACG | IBCh | 2.54±0.60 | 0.60±0.12 | 2.00±0.23 | -0.54±0.64 |
| GCAUU1GC<br>CG1UUACG | IBCh | 2.19±0.61 | 0.60±0.12 | 2.00±0.23 | -0.19±0.65 |
| GGAC11CUGG<br>CCUGCCGACC | DS | 2.59±0.45 | 1.21±0.19 | 1.21±0.19 | -1.39±0.49 |
| GGAC11GUCC<br>CCUG11CAGG | DS | -1.32±0.59 | -0.44±0.23 | -2.3±0.34 | -0.98±0.68 |
| GGAG11CUCC<br>CCUC11GAGG | DS | -2.34±0.61 | -0.57±0.31 | -2.43±0.40 | -0.09±0.73 |
| GCG1U1CGC | IBCh | -1.86±0.48 | 0.83±0.42 | -0.96±1.07 | 0.90±1.18 |

| | | | | | |
|---|---|---|---|---|---|
| CGC1U1GAG | | | | | |
| GCG1UUCGC<br>CGCUU1GCG | IBCh | -2.58±0.50 | 0.83±0.42 | -1.92±1.09 | 0.66±1.20 |
| GCGU1UCGC<br>CGCU1UGCG | IBCh | 0.05±0.43 | 0.83±0.42 | 0.90±1.04 | 0.85±1.13 |

Table S15: Internal loop-contained duplexes studied by optical melting to test the internal loop parameters. The character "1" is used to represent 1mΨ. These experiments were performed at DNA Software, Inc.

| | Average of curve fits | | | | $T_M^{-1}$ vs log $C_T$ plots | | | |
|---|---|---|---|---|---|---|---|---|
| Sequence | -ΔH° (kcal/mol) | -ΔS° (eu) | -ΔG°37 (kcal/mol) | $T_M$† (°C) | -ΔH° (kcal/mol) | -ΔS° (eu) | -ΔG°37 (kcal/mol) | $T_M$† (°C) |
| CGCCAGAGCCGG<br>GCG11CUC1GCC | 94.92±6.29 | 264.92±19.16 | 12.75±0.36 | 58.8 | 97.02±14.19 | 271.37±43.48 | 12.85±0.77 | 58.6 |
| CGC1AGAG1CGG<br>GCGC1CUCCGCC | 89.99±3.38 | 249.01±10.3 | 12.76±0.2 | 60.1 | 92.3±7.39 | 256.11±22.68 | 12.86±0.38 | 59.9 |
| GACCGAGACCAC<br>CUG1CUC11GUG | 97.01±5.77 | 273.86±17.82 | 12.07±0.27 | 55.8 | 88.13±8.41 | 246.4±26 | 11.71±0.38 | 56.4 |
| GGC1GAGA1CGC<br>CCGCCUC1CGCG | 109.47±9.06 | 305.52±27.23 | 14.72±0.63 | 62.1 | 110.58±17.49 | 308.96±52.97 | 14.76±1.12 | 61.9 |

†$T_M$ is calculated for strand concentration of 0.1 mM.

Table S16: Agreement between internal loop experiments and nearest neighbor estimates of $\Delta G°_{37}$. This shows the duplex stabilities ($\Delta G°_{37}$ Measured), the estimated stabilities ($\Delta G°_{37}$ Estimated), and their differences ($\Delta\Delta G°_{37}$).

| Sequence | $\Delta G°_{37}$ Measured (kcal/mol) | $\Delta G°_{37}$ Estimated (kcal/mol) | $\Delta\Delta G°_{37}$ (kcal/mol) |
|---|---|---|---|
| CGCCAGAGCCGG<br>GCG11CUC1GCC | -12.85±0.51 | -13.01±0.84 | 0.16±0.98 |
| CGC1AGAG1CGG<br>GCGC1CUCCGCC | -12.86±0.51 | -13.01±0.84 | 0.15±0.98 |
| GACCGAGACCAC<br>CUG1CUC11GUG | -11.71±0.47 | -10.86±0.83 | -0.85±0.95 |
| GGC1GAGA1CGC<br>CCGCCUC1CGCG | -14.76±0.59 | -14.38±0.84 | -0.38±1.02 |

Table S17. Optical melting data for duplexes containing bulge loops. The character "1" is used to represent 1mΨ. Site is either the Institute of Bioorganic Chemistry (IBCh) or DNA Software (DS).

| | | Average of curve fits | | | | $T_M^{-1}$ vs log $C_T$ plots | | | |
|---|---|---|---|---|---|---|---|---|---|
| Sequence | Site | -ΔH° (kcal/mol) | -ΔS° (eu) | -ΔG°37 (kcal/mol) | $T_M$† (°C) | -ΔH° (kcal/mol) | -ΔS° (eu) | -ΔG°37 (kcal/mol) | $T_M$† (°C) |
| CCA-AAGCA<br>AGG1G1UCG | DS | 87.2±11.0 | 252.6±35.2 | 8.89±0.17 | 45.6 | 77.7±9.2 | 222.0±29.4 | 8.83±0.25 | 46.5 |
| GCG1GCG<br>CGC-CGC | IBCh | 62.8±3.9 | 172.6±12.1 | 9.28±0.15 | 51.1 | 66.3±2.3 | 183.6±7.1 | 9.39±0.08 | 50.8 |
| GACC1UGUC<br>CUGG-ACAG | IBCh | 64.7±3.8 | 179.2±11.8 | 9.18±0.16 | 49.9 | 62.5±2.8 | 172.3±8.7 | 9.09±0.08 | 50.1 |
| UGAC1CUCA<br>ACUG-GAGU | IBCh | 61.8±4.9 | 172.4±15.7 | 8.36±0.17 | 46.3 | 63.2±3.9 | 176.9±12.4 | 8.37±0.09 | 46.1 |

†$T_M$ is calculated for strand concentration of 0.1 mM.

Table S18. Agreement between experimental and nearest neighbor estimated $\Delta G°_{37}$ for bulge loops. The predicted stability is the free energy cost of a single bulge and is sequence independent [15]. Therefore, the U analog stability is the nearest neighbor prediction.

| Sequence: | $\Delta G°_{37\ loop}$ (kcal/mol) | $\Delta G°_{37\ loop}$ (U analog/Prediction) (kcal/mol) | $\Delta\Delta G°_{37\ loop}$ (Prediction-Measured) (kcal/mol) |
|---|---|---|---|
| CCA-AAGCA<br>AGG1G1UCG | 3.65±0.70 | 3.89±0.62 | 0.24±0.94 |
| GCG1GCG<br>CGC-CGC | 1.34±0.50 | 3.89±0.62 | 2.55±0.80 |
| GACC1UGUC<br>CUGG-ACAG | 4.06±0.58 | 3.89±0.62 | -0.17±0.85 |
| UGAC1CUCA<br>ACUG-GAGU | 3.14±0.46 | 3.89±0.62 | 0.75±0.77 |

Table S19. tRNA structure prediction accuracies for U and 1mΨ. Ensemble Defect U is the normalized ensemble defect for folding to the native cloverleaf structure when U is used for 1mΨ in the sequence. Ensemble Defect 1mΨ is the ensemble defect when 1mΨ is used in the sequence. For ensemble defect, lower values represent modeling that is more consistent with the known cloverleaf structure. MEA Sensitivity U is the percent of known pairs correctly predicted using maximum expected accuracy structure prediction when U is used in the sequence in place of 1mΨ. MEA Sensitivity 1mΨ is the percent of known pairs correctly predicted using maximum expected accuracy structure prediction when 1mΨ is used in the sequence. MEA PPV U is the percent of predicted pairs that are in the known cloverleaf structure using maximum expected accuracy structure prediction when U is used in the sequence in place of 1mΨ. MEA PPV 1mΨ is the percent of predicted pairs that are in the known cloverleaf structure using maximum expected accuracy structure prediction when 1mΨ is used in the sequence. For Sensitivity and PPV (Positive Predictive Value), higher numbers are better agreement between the modeling and the known cloverleaf structure.

| tRNA Accession | Ensemble Defect U | Ensemble Defect 1mΨ | MEA Sensitivity U | MEA Sensitivity 1mΨ | MEA PPV U | MEA PPV 1mΨ |
|---|---|---|---|---|---|---|
| RA0380 | 0.198 | 0.190 | 100.0 | 100.0 | 91.3 | 91.3 |
| RA0500 | 0.263 | 0.194 | 95.2 | 100.0 | 87.0 | 91.3 |
| RA0501 | 0.310 | 0.274 | 76.2 | 81.0 | 80.0 | 100.0 |
| RA0502 | 0.098 | 0.078 | 95.2 | 95.2 | 90.9 | 90.9 |
| RC0500 | 0.210 | 0.157 | 89.5 | 89.5 | 81.0 | 81.0 |
| RD0500 | 0.472 | 0.424 | 63.2 | 89.5 | 60.0 | 85.0 |
| RE0500 | 0.238 | 0.235 | 89.5 | 89.5 | 89.5 | 89.5 |
| RE0501 | 0.560 | 0.545 | 63.2 | 63.2 | 54.6 | 54.6 |
| RF0500 | 0.419 | 0.426 | 100.0 | 100.0 | 100.0 | 100.0 |
| RG0380 | 0.132 | 0.125 | 85.0 | 85.0 | 81.0 | 81.0 |
| RG0500 | 0.076 | 0.061 | 100.0 | 100.0 | 95.5 | 95.5 |
| RG0501 | 0.132 | 0.125 | 85.0 | 85.0 | 81.0 | 81.0 |
| RG0502 | 0.132 | 0.125 | 85.0 | 85.0 | 81.0 | 81.0 |
| RG0503 | 0.086 | 0.072 | 100.0 | 100.0 | 100.0 | 100.0 |
| RG4800 | 0.453 | 0.448 | 65.0 | 65.0 | 86.7 | 86.7 |
| RH0380 | 0.320 | 0.304 | 100.0 | 100.0 | 73.1 | 73.1 |
| RI0500 | 0.218 | 0.216 | 95.2 | 95.2 | 90.9 | 90.9 |
| RI0501 | 0.288 | 0.300 | 57.1 | 57.1 | 46.2 | 44.4 |
| RK0500 | 0.341 | 0.309 | 81.0 | 81.0 | 73.9 | 73.9 |
| RK0501 | 0.304 | 0.311 | 76.2 | 76.2 | 80.0 | 72.7 |
| RL0500 | 0.278 | 0.247 | 95.5 | 95.5 | 100.0 | 100.0 |
| RL0501 | 0.250 | 0.245 | 82.6 | 82.6 | 79.2 | 79.2 |
| RL0502 | 0.310 | 0.309 | 87.0 | 87.0 | 83.3 | 83.3 |
| RL0503 | 0.248 | 0.193 | 87.0 | 87.0 | 76.9 | 76.9 |
| RL0504 | 0.235 | 0.250 | 87.0 | 87.0 | 87.0 | 87.0 |

| | | | | | | |
|---|---|---|---|---|---|---|
| RM0500 | 0.477 | 0.471 | 60.0 | 60.0 | 46.2 | 46.2 |
| RN0380 | 0.137 | 0.113 | 100.0 | 95.2 | 95.5 | 95.2 |
| RN0500 | 0.037 | 0.024 | 100.0 | 100.0 | 100.0 | 100.0 |
| RP0500 | 0.480 | 0.479 | 36.8 | 36.8 | 29.2 | 29.2 |
| RP0501 | 0.279 | 0.243 | 89.5 | 100.0 | 81.0 | 79.2 |
| RP0502 | 0.417 | 0.373 | 89.5 | 89.5 | 100.0 | 100.0 |
| RR0380 | 0.261 | 0.261 | 100.0 | 100.0 | 76.0 | 76.0 |
| RR0500 | 0.100 | 0.086 | 100.0 | 100.0 | 95.5 | 95.5 |
| RR0501 | 0.317 | 0.330 | 100.0 | 100.0 | 76.0 | 76.0 |
| RR0502 | 0.523 | 0.544 | 81.0 | 95.2 | 94.4 | 95.2 |
| RS0380 | 0.137 | 0.137 | 100.0 | 100.0 | 92.0 | 92.0 |
| RS0500 | 0.276 | 0.224 | 82.6 | 87.0 | 73.1 | 76.9 |
| RS0501 | 0.487 | 0.473 | 100.0 | 100.0 | 88.9 | 88.9 |
| RS0502 | 0.324 | 0.311 | 100.0 | 100.0 | 92.0 | 92.0 |
| RT0380 | 0.101 | 0.102 | 100.0 | 100.0 | 91.3 | 91.3 |
| RT0500 | 0.058 | 0.055 | 100.0 | 100.0 | 91.3 | 91.3 |
| RT0501 | 0.106 | 0.101 | 100.0 | 100.0 | 91.3 | 91.3 |
| RV0380 | 0.454 | 0.462 | 60.0 | 85.0 | 50.0 | 77.3 |
| RV0381 | 0.359 | 0.359 | 75.0 | 75.0 | 55.6 | 55.6 |
| RV0382 | 0.340 | 0.340 | 60.0 | 60.0 | 48.0 | 48.0 |
| RV0500 | 0.162 | 0.162 | 90.0 | 90.0 | 100.0 | 100.0 |
| RV0501 | 0.260 | 0.257 | 60.0 | 60.0 | 54.6 | 54.6 |
| RW0500 | 0.314 | 0.307 | 76.2 | 71.4 | 80.0 | 71.4 |
| RW8580 | 0.148 | 0.110 | 100.0 | 100.0 | 100.0 | 100.0 |
| RX0380 | 0.322 | 0.255 | 95.0 | 95.0 | 100.0 | 100.0 |
| RX0500 | 0.303 | 0.235 | 80.0 | 80.0 | 76.2 | 76.2 |
| RX0540 | 0.218 | 0.221 | 85.0 | 85.0 | 77.3 | 77.3 |
| RY0500 | 0.095 | 0.093 | 81.0 | 81.0 | 73.9 | 73.9 |
| RY9990 | 0.444 | 0.441 | 95.0 | 95.0 | 100.0 | 100.0 |
| RY9991 | 0.152 | 0.110 | 95.0 | 95.0 | 100.0 | 100.0 |

Table S20. Agreement between prior stacking parameters and these current parameters.

| Stack: | Prior: $\Delta G^\circ_{37}$ (kcal/mol) | Current: $\Delta G^\circ_{37}$ (kcal/mol) |
|---|---|---|
| G1<br>CA | -2.43±0.08 | -2.80±0.24 |
| 1G<br>AC | -2.26±0.07 | -2.58±0.25 |
| 1C<br>AG | -2.67±0.08 | -2.53±0.25 |
| C1<br>GA | -1.83±0.09 | -2.22±0.24 |
| 1A<br>A1 | -1.86±0.15 | -1.81±0.45 |
| A1<br>1A | -1.13±0.12 | -1.51±0.46 |
| 11<br>AA | -1.18±0.40 | -1.23±0.25 |
